# Biotic modulators of global change effects on plant communities

**DOI:** 10.64898/2026.02.11.704804

**Authors:** Anu Eskelinen, Martin Andrzejak, W. Stanley Harpole, Susan Harrison, Aimée T. Classen, Anna-Liisa Laine, Noémie A. Pichon, Anita C. Risch, Jake Alexander, Maria-Theresa Jessen, Phoebe L. Zarnetske, Lotte Korell

## Abstract

Understanding and predicting future plant biodiversity and productivity is critical for prioritizing global change mitigation, conservation, and restoration efforts. One major challenge is that we know remarkably little of how interspecific interactions may modulate the effects of global change factors on diversity and productivity. Here, we develop and test a synthetic conceptual framework about how different ‘biotic modulators’ (herbivory, plant-plant interactions, pathogens, mycorrhiza) can either amplify or mitigate the effects of global change drivers (nutrient and CO_2_ enrichment, changes in rainfall and temperature) on plant community biomass and diversity. We report that herbivores mitigated both biomass increment and diversity decline caused by different global change drivers, while plant competition did not significantly alter global change impacts due to mixed effects (both amplification and mitigation). Pathogens tended to function similarly to herbivores, while mycorrhiza both amplified and mitigated community responses. Our conceptual framework further identifies mechanisms by which species interactions can modify global change effects, provides new testable hypotheses, and identifies research gaps and future research directions. We conclude that plant consumers can be important agents stabilizing plant productivity and safeguarding plant biodiversity in the Anthropocene, while more research is urgently needed to understand the role of other biotic modulators.

## I. INTRODUCTION

Generalizing global change effects on plant biodiversity and productivity has proven challenging because idiosyncratic and highly context-dependent responses of local communities to different global change drivers are often observed (Dornelas *et al*., 2014; Vellend *et al*., 2017). For example, while declines in plant biodiversity have been shown in response to some global change drivers (Harrison, Gornish & Copeland, 2015; Vellend *et al*., 2017; Jandt *et al*., 2022), stable or increasing richness combined with altered community composition are seen in other cases (Vellend *et al*., 2017; Steinbauer *et al*., 2018; Nielsen *et al*., 2019; Pilotto *et al*., 2020). The major challenges and opportunities for better understanding and predicting this high context-dependency rely on identifying which factors and conditions in local communities cause these heterogeneous responses to global change drivers.

We propose that ‘biotic modulators’ of global change effects, i.e., interactions among plant species and other organisms, could provide a key to elucidate these heterogeneous responses to global change drivers (Fig. 1). Some previous individual studies indicate that plant competitive or facilitative interactions, plant-consumer interactions with herbivores and pathogens, and plant-mycorrhizal interactions can be critical in determining how different global change factors manifest themselves in local plant communities (Alexander *et al*., 2015; Classen *et al*., 2015; Kaarlejärvi *et al*., 2017; Vandvik *et al*., 2020; Post *et al*., 2023). Although the role of biotic modulators has been acknowledged in general (Tylianakis *et al*., 2008; Zarnetske *et al*., 2012; Urban *et al*., 2016), none of the previous conceptual papers have attempted to quantitatively synthesize what generalities emerge from existing data and, importantly, none of them focuses on plant biodiversity and productivity, which are essential to sustaining life, biodiversity, and ecosystem functions on earth. Knowledge on how different modulators alter plant biodiversity and productivity responses to global change drivers is therefore scant, and we lack a synthesis of what is already known.

**Figure 1.**
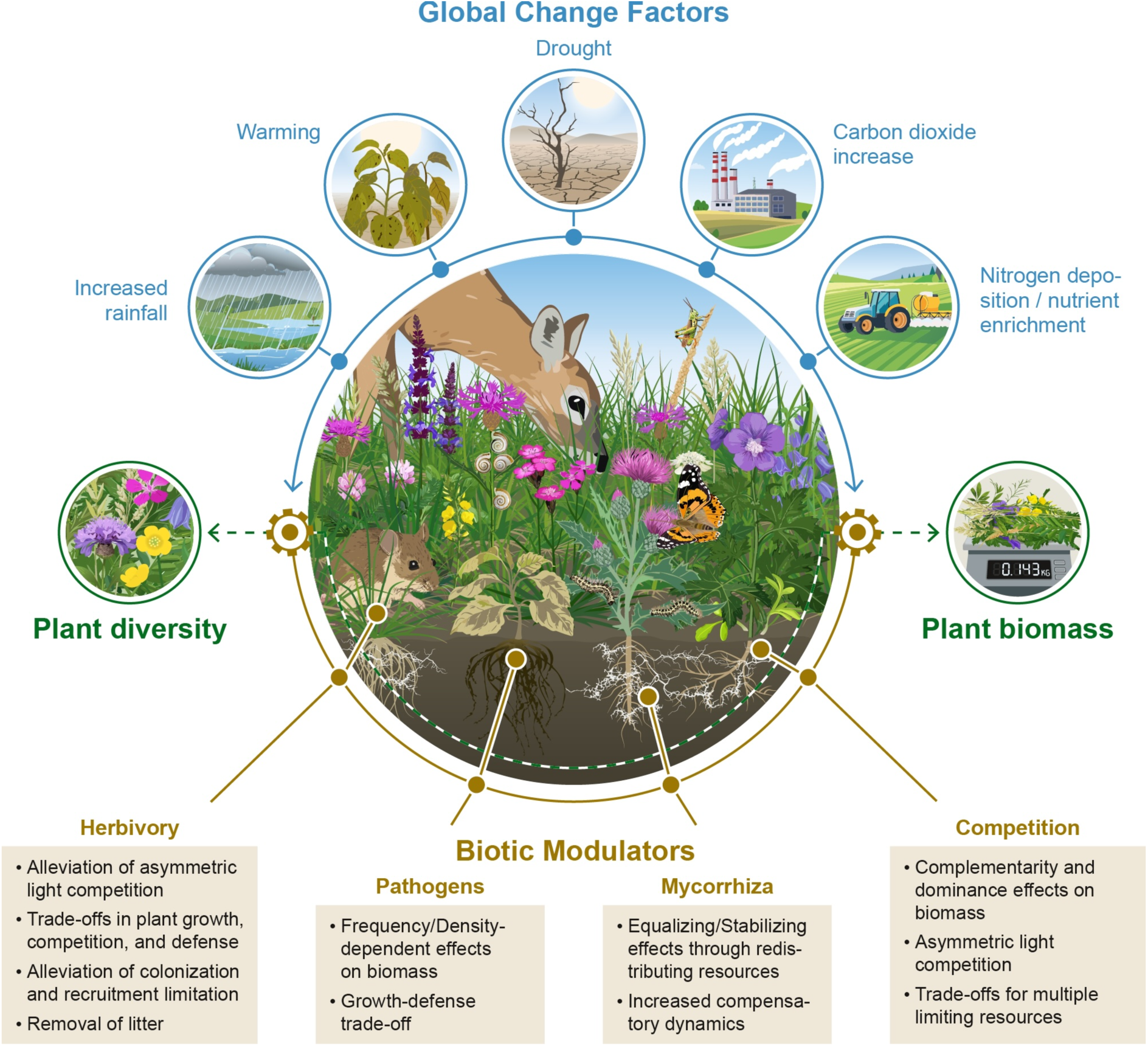
A conceptual illustration of how different biotic factors (herbivory, pathogens, mycorrhiza, plant competition; hence ‘biotic modulators’) can alter the effects of global change factors (increased rainfall, warming, drought, increased CO_2_, and nutrient enrichment) on biomass production and diversity of terrestrial plant communities. Biotic modulators can either amplify or mitigate global change effects, or have neutral effects. Besides interacting with global change factors, different modulators could, in theory, also exhibit complex interactions with each other; these interactions are not illustrated for clarity. For example, the effects of some modulators (e.g., herbivory and pathogens) on plant community biomass and diversity operate through altered plant competitive interactions. Under each modulator, we list mechanisms th<u>r</u>ough which the modulator can alter global change effects; these are discussed in more detail in the text.

Many global change factors directly or indirectly alter the availability of essential resources to plant growth (i.e., nutrients, water, light) and therefore possess the capacity to fundamentally alter plant productivity and all biotic interactions that depend on plant resource-use and biomass production. For example, nutrient enrichment, increased precipitation and drought can alter plant competitive outcomes, as different species are likely the best competitors in low vs. high resource availability (Tilman, 1982; Craine & Dybzinski, 2013; Van Dyke, Levine & Kraft, 2022). While temperature is not a resource per se, it can indirectly augment soil nutrient availability through enhanced microbial activity and decomposition in warming soils (Doetterl *et al*., 2022), therefore altering species interactions (Alexander *et al*., 2015). Further, any changes in plant communities should also reflect on to plant consumers, such as herbivores and pathogens, or symbionts, such as mycorrhiza, that depend on primary producers and can also impose strong control over both plant community biomass and diversity (Van Der Heijden *et al*., 1998; Mitchell *et al*., 2003; Berver *et al*., 2015; Kaarlejärvi *et al*., 2017; Borer *et al*., 2020; Paseka *et al*., 2020; Haman *et al*., 2021; Eskelinen *et al*., 2022; Post *et al*., 2023). These within and between trophic level interactions may mitigate or amplify (i.e., lessen or increase) productivity and diversity responses to global change drivers.

We developed a novel conceptual framework of biotic modulators to increase the understanding of how and by which mechanisms species interactions within and between trophic levels can alter global change effects on plant communities (Fig. 1). We identify when and why different modulators might amplify or mitigate the effects on how different global change factors affect plant diversity and biomass, how the nature of different global change factors (e.g., either resource-reducing or -enhancing) affects its interaction with the modulators, what are the underlying mechanisms, for example, resource competition, trade-offs, and complementarity/dominance effects, by which modulators alter global change effects on plant communities (Fig. 1); these aspects of the framework are discussed in detail in the discussion. To test our framework, we conducted a systematic literature review to synthesize what is known about biotic modulators in mitigating or amplifying global change effects on plant community biomass and diversity. We reviewed 8789 abstracts and identified 51 experimental studies that manipulated both a global change factor (nutrients, water, temperature, CO_2_) and a biotic modulator (herbivory, pathogens, mycorrhiza, plant-plant interactions), and reported results on total community biomass and diversity. Individual species and population-level responses were omitted as whole community responses cannot be inferred from them. We chose these global change drivers as they alter resource availability to plants either directly or indirectly, and can therefore alter species interactions. We chose these modulators as they can be manipulated on plot-level scale where global change manipulations and plant community studies are conducted. Although our literature review/synthesis was not limited to any particular terrestrial systems, experimental research in this area is dominated by especially grassland ecosystems. We conducted a meta-analysis to synthesize existing data for herbivory and plant competition, as the study number allowed a quantitative analysis for these two modulators. For pathogens and mycorrhiza, we conducted a numerical review of all cases that fit the search criteria, as there were fewer studies. Based on the meta-analyses and review, our conceptual framework uncovers mechanisms of biotic modulation and identifies generalizations, critical knowledge gaps, and future research directions (Fig. 1).

## II. METHODS

### (1) Choice of studies

We focused on interactions between global change factors and biotic modulators and did not include additive or main effects as our inherent interest is in how different biotic modulators alter the effects of global change factors on plant communities. We were also not interested in the effects of either global change or biotic factors alone. These have been addressed in multiple previous studies (synthesized by Wu *et al*., 2011; Komatsu *et al*., 2019; Korell *et al*., 2021) and are out of the scope of this paper. We also did not include biodiversity manipulation studies in our literature review. Further, we only focused on experimental studies because we wanted to assure there are causal links between the global change factors, biotic modulators, and the observed changes in community-level plant biomass and diversity/richness. We acknowledge the value of studies carried out along environmental gradients (e.g. observational studies, manipulation of biotic modulators along environmental gradients) but since many factors can change along gradients in general and this would diminish our ability to infer causality between the studied variables, we decided to focus on experimental studies only. When a single experiment reported several response variables, for example, both community biomass and diversity, we included them all to our review. The number of separate studies has been reported in Tables S1-S4.

***Global change drivers*** In our global change search, we included nutrient enrichment, climate warming, changes in precipitation (both increased and decreased precipitation/drought), and CO_2_ increase. Although habitat loss and fragmentation are equally important global change drivers, we did not include them in our list of drivers because they often operate on larger scales, are difficult to manipulate experimentally and do not directly alter resource availability to plants. The global change factors that we chose can all be experimentally manipulated on a plot-level in natural systems, and all directly alter resource availability to plants, either nutrient, water or CO_2_ availability, and are therefore likely to change plant productivity and interactions among plants (competition, facilitation), their consumers (herbivory, plant-pathogen interactions), and mutualists (mycorrhizal interactions).

***Biotic modulators*** We considered plant-plant interactions (competition and facilitation), aboveground herbivory (both insect and mammalian), pathogens, and mycorrhizal fungi as biotic modulators. We did not include pollinators as a biotic modulator group in the study because their effects are harder to study on a small plot-level scale where global change manipulations and plant community studies are conducted.

***Plant community measures*** We chose total aboveground plant community biomass as our community level surrogate for abundance/productivity but also accepted total plant cover and biomass/abundance estimates obtained through using point intercept method, if biomass measurements were not available. This was considered important as collecting biomass is destructive for vegetation and often not possible to collect in long-term experiments. For diversity measures, we included species richness and different diversity indexes (such as Shannon and Simpson diversity), and also included studies reporting seedling richness and/or diversity. If individual studies reported both species richness and diversity, we included richness and one diversity index (as richness and diversity indexes can show different responses); therefore, our data may include maximum two diversity measures from one study. For consistency, we did not include evenness among our diversity measures as it was rarely measured. Further, the studies reporting evenness usually reported also richness and Shannon/Simpson diversity, and evenness describes a slightly different aspect of diversity that is also included in Shannon and Simpson diversity indeces.

As our focus was on community-level changes in response to interactions between global change factors and biotic modulators, we did not include studies investigating individual plant, functional group or other species group level responses. We are aware that especially studies of plant competition and plant-mycorrhizal interactions are often on individual species level, so that results are reported for individual plant or species performance. However, individual species or functional group responses do not reflect whole community responses (neither biomass nor richness) and could be opposite to the responses of other species/functional groups; we therefore excluded these studies.

We did not restrict our studies on herbaceous plant communities, i.e., meadows and grasslands, and our search can therefore include also shrublands and forests. However, as forest ecosystems are harder to experimentally manipulate, most studies originate from herbaceous systems, in particular grassland ecosystems (Tables S1-S4).

### (2) Literature search

To investigate how biotic modulators alter the effects global change factors on plant community biomass and diversity we conducted a systematic literature review. We conducted our initial search in the Web of Science search engine separately for each biotic modulator. In general, since our search criteria were strict, i.e., each study had to experimentally manipulate both a global change factor and a biotic modulator, and report results of their interactions on community biomass and/or richness/diversity, our initial screening included a lot of search terms and produced a long list of cases (herbivory, 447 papers; competition, 5763 papers; pathogens, 1818 papers; mycorrhiza, 761 papers), while after the manual screening the list of relevant cases was usually very short (Tables S1-S4). We complemented the search in Web of Science by manual screening of papers that were cited in the reference lists of the relevant papers and that were citing these papers, and found some additional studies; the final data set included 51 primary studies (herbivory, 26 papers; competition, 14 papers; pathogens, 5 papers; mycorrhiza, 6 papers; see Figs. S2-S5 for Prisma charts). The phrases used in the in the initial literature search for each biotic modulator are reported in Supplementary File.

### (3) Statistical analyses

***Data extraction and interpretation*** For each individual study found to be relevant in the systematic literature search, we extracted the mean and standard errors (SE) from the relevant figures using the ‘metaDigitise’ package in R (Pick, Nakagawa & Noble, 2019). In one case where neither standard errors nor standard deviations were reported in the article, we contacted the author of the study and calculated the needed statistics ourselves. Also, in one case we extracted data from a figure that was unpublished but which we knew about (because one of us authored the paper). We then calculated log response ratio (LRR) as an effect size for the comparisons of interest. First, we defined the direct impact of the global change driver (LRR_GC_ = Ln(effect of global change factor in the absence of the biotic modulator / control in the absence of the biotic modulator) and assessed whether this effect was neutral, positive or negative (i.e., whether it had no effect, increased or decreased diversity and/or biomass). Second, we classified the effect of the biotic modulator by comparing the impact of the global change factor in the absence vs. in the presence of the modulator (LRR_BM_ = Ln(effect of the global change factor in the absence of the biotic modulator / effect of the global change factor in the presence of the biotic modulator). Biotic modulator could either mitigate (make the impact smaller) or amplify (make the impact greater) the effect of the global change factor, have a neutral effect. For the meta-analyses we also calculated the sample variance corresponding to the LRR based on the standard deviation of the primary study (either given directly or calculated by using the standard error).

***Analyses*** We found enough studies for quantitative meta-analyses of herbivory and competition (26 and 14 studies, respectively). We performed these meta-analyses separately as these were also separate search terms in our systematic literature search. We had too few studies for pathogens and mycorrhiza to allow meta-analyses and we therefore conducted a numerical review of all cases that fit the search criteria. For pathogens and mycorrhiza, we refer to changes in effect sizes that we calculated from individual studies when assessing the significance of primary studies that we reviewed.

All analyses were performed using R statistical environment (version 4.4.1; R Core Team, 2024). We used linear mixed effects meta-regression models in ‘metafor’ package in R (Viechtbauer, 2010) to conduct the meta-analyses. We separately analyzed the responses of diversity measures (e.g., species richness, Shannon diversity) and productivity measures (e.g. biomass, aboveground net primary productivity (ANPP)). In all models, fixed predictor variables were the type of manipulation (sole effect of global change factor or the impact of modulator) and the type of global change factor (nutrient enrichment, climate warming, increased precipitation, drought, and CO_2_ increase). As some studies included multiple global changes drivers and/or multiple diversity metrics, we included study as a random factor in all models. Further, the models were weighted by the corresponding sample variance of LRR, to account for the variability in the results of each primary study.

We did not include interaction between the type of manipulation and the type of global change factor to avoid overparameterizing the models, as the sample size (i.e., number of studies) was low for both herbivory and competition. We also did not separately study the impact of the type of herbivory (vertebrate, invertebrate, or both) for the same reasons.

To test for publication bias (Sterne & Egger, 2001), we used funnel plots via the ‘funnel’ function of the ‘metafor’ package (Viechtbauer, 2010). For the modulators analyzed quantitatively, we detected significant signs of publication bias (Fig. S1), which is a common finding in meta-analyses (e.g.,Trepel *et al*., 2024).

## III. RESULTS

### (1) General

For herbivory, most studies manipulated vertebrate herbivory using different-sized exclosures to exclude herbivores, while a minority manipulated invertebrate herbivory using insecticide (Table S1). For plant competition, studies used variable methods to reduce competition, including functional group, random or dominant plant removals, tiebacks (competitors pulled aside using a net), light addition (to relax competition for light), and also total biomass removal or disturbance when the response variable was seedling richness/diversity (Table S2). For pathogens, the studies used mostly foliar fungicide to reduce pathogens (Table S3). For mycorrhiza, the studies used Benomyl fungicide specific to arbuscular mycorrhiza (AM) to reduce AM, or communities were grown completely without AM (mesocosm studies; Table S4).

The majority of the studies across all modulator groups manipulated nutrients, mimicking nutrient enrichment as the global change driver (Tables S1-S4). While warming was the next commonly manipulated global change driver, there were very few studies addressing interactions among a biotic modulator, precipitation increase, drought, and CO_2_ enrichment.

### (2) Herbivory

***Biomass*** We found that the herbivory significantly altered the impact of global change factors on plant community biomass (Figs. 2a and 3a, Table S5). Overall, global change factors enhanced plant community biomass in the absence of herbivory and herbivory mitigated this increase (11 cases; 8 nutrients, 1 increased precipitation, 1 warming, 1 nutrients and warming; Figs. 2a, 3a, Table S1). In 2 cases, nutrients increased biomass and herbivores amplified this increase, while in 1 case, drought decreased biomass and herbivores amplified this negative effect (Fig. 2a, Table S1). The type of global change factor did not significantly affect this outcome (Fig. 3a, Table S5); however, we had very few studies manipulating precipitation (2) and temperature (7), and no studies addressing interaction between CO_2_ and herbivory (Table S1).

**Figure 2.**
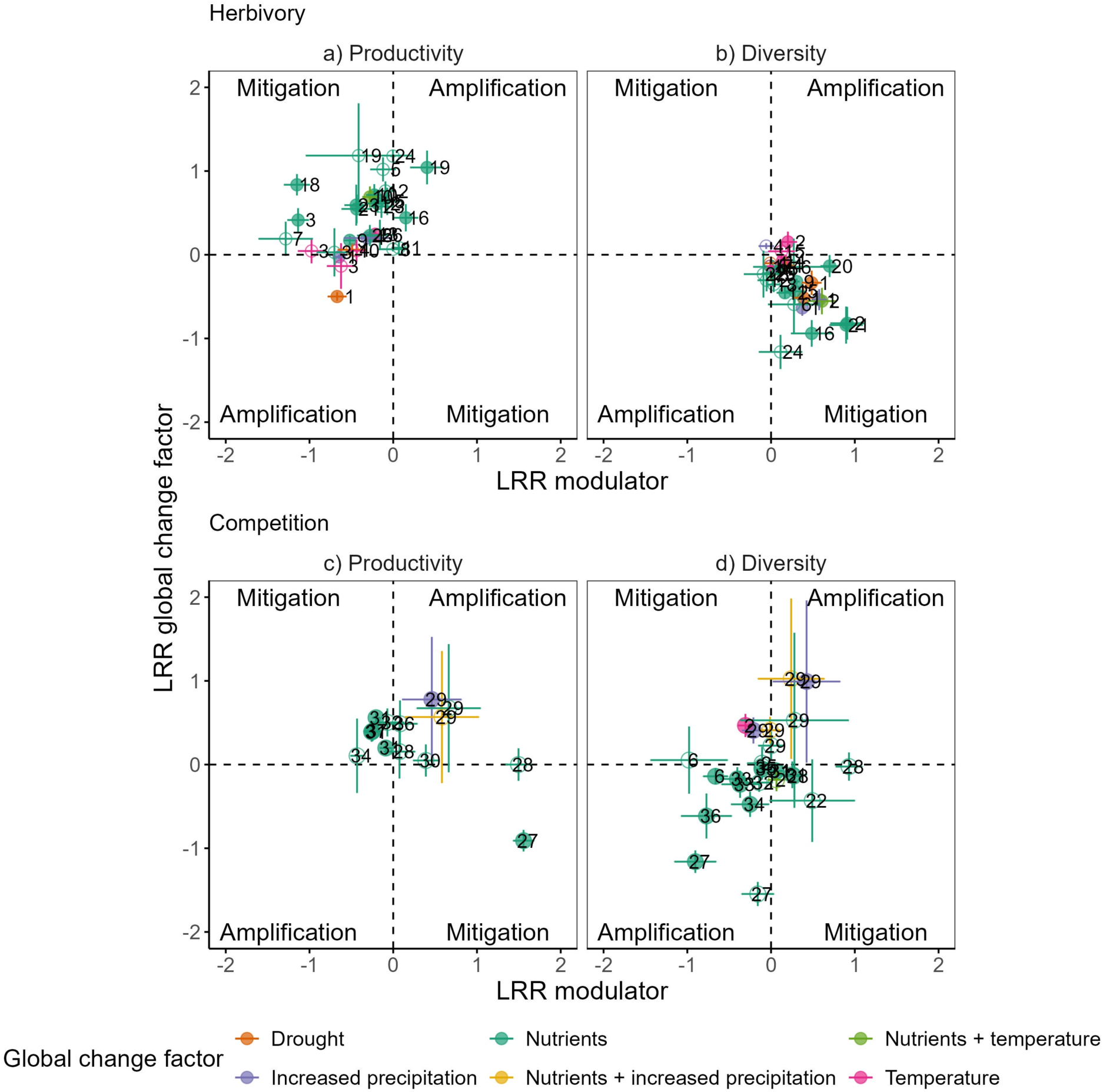
Dependence of the log response ratio (LRR) of diversity (a,c) and productivity (b,d) for the different global change factors (CO2, increased precipitation, drought, nutrient increase, temperature, nutrients + temperature, nutrients + increased precipitation) and the LRR of the biotic modulators herbivory (a,b) and competition (c,d). Positive values for LRR of the global change factor (y-axis) indicate a positive effect of the global change on diversity (a,c) or productivity (b,d) in the absence of the biotic modulators herbivory or competition relative to the control in the absence of the herbivory or competition, whereas negative values indicate a negative effect of the global change factor on diversity (a,c) or productivity (b,d) in the absence of the biotic modulators herbivory or competition relative to the control. As the LRR of the biotic modulator describes the impact of the global change factor in the absence relative to the presence of the modulator (herbivory or competition), mitigation (make the impact smaller) or amplification (make the impact greater) could be either indicated by positive or negative LRR values as they depend on the effect of the global change factor on diversity and productivity. Different corners of the diagram indicate the direction of biotic modulation (mitigation, amplification). Filled circles indicate that the LRR and their SE do not overlap with zero, whereas open circles indicate an overlap with zero, i.e., neutral effect.

**Figure 3.**
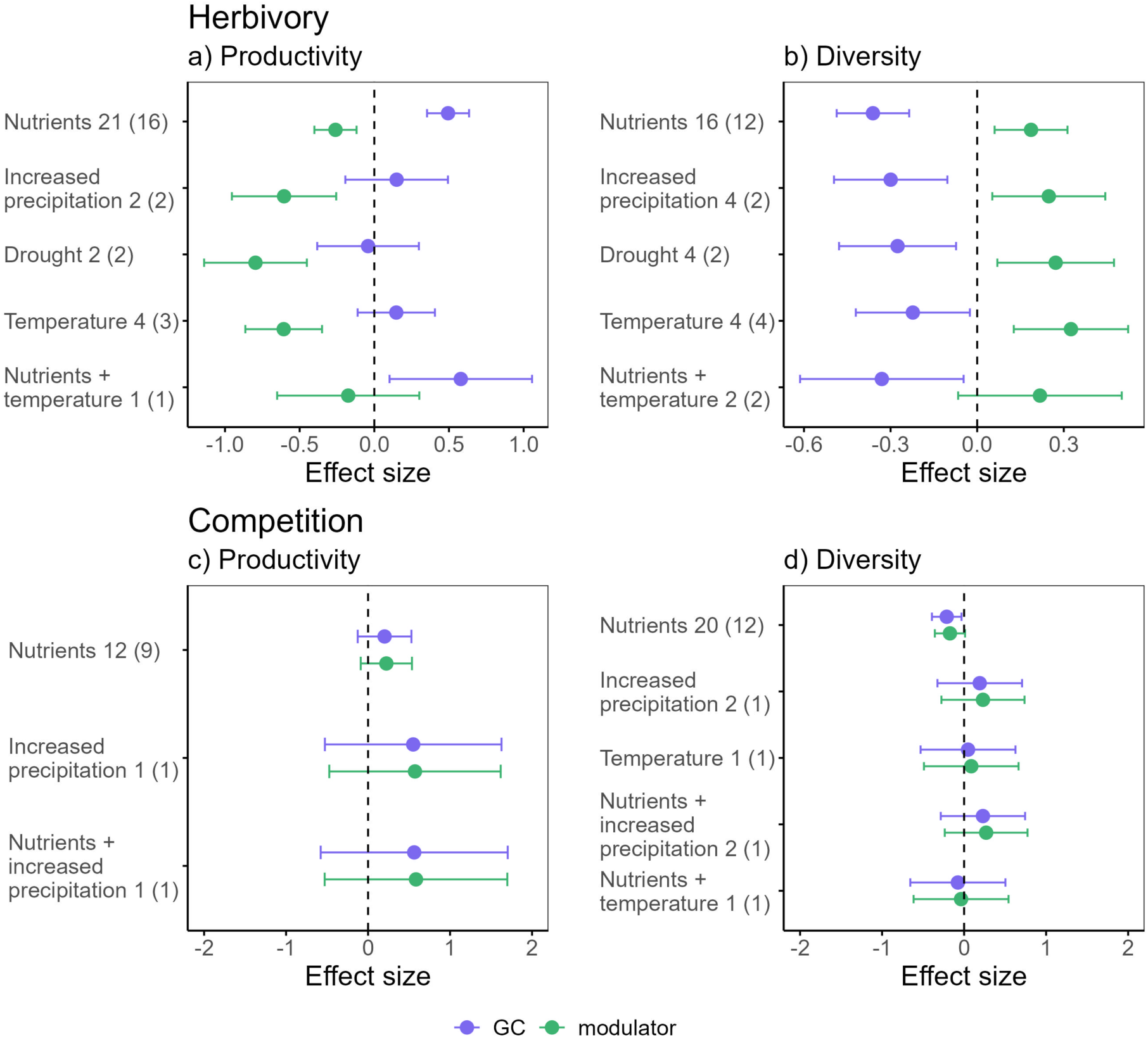
Overall effect size estimates for productivity (a,c) and diversity (b,d) and upper and lower confidence interval extracted from metanalytical model. Estimates are given for the sole effect of the respective global change driver (GC, violet) and when the modulators herbivory (a,b) or competition (c,d) are present (modulator, green) separately for the different global change factors (increased precipitation, drought, nutrient increase, temperature, nutrients + temperature, nutrients + increased precipitation). Small numbers next to the global change factor indicate the number of effect sizes included in the model where the first number corresponds to the overall number of cases included and the number in parentheses gives the number of the respective studies.

***Diversity*** Similarly, we found that herbivory significantly modified the impact of global change factors on plant community diversity (Figs. 2b and 3b, Table S5). Overall, global change factors decreased plant diversity and herbivory mitigated this decline (15 cases; 8 nutrients, 3 drought, 2 increased precipitation, 1 warming, 1 nutrients and warming; Figs. 2b and 3b, Table S1). Again, the type of global change factor had no impact (Fig. 3b, Table S5).

### (3) Plant competition

***Biomass*** We found that, overall, plant competition did not significantly modify the impact of the global change factors on plant biomass; type of global change factor also showed no significant effect (Figs. 2c and 3c, Table S6). Individual studies represented a mixture of amplification, mitigation and neutral responses. For example, in 4 cases, nutrients increased biomass and competition mitigated this increase, i.e., there was smaller effect on biomass when competition was not reduced. In contrast, in 1 case, competition amplified a positive biomass response to precipitation increase (Fig. 2c, Table S2). In another case, nutrients decreased biomass and competition mitigated this increase. There were several neutral cases (Fig. 2c, Table S2), and no studies manipulating drought, temperature and CO_2_.

***Diversity*** Similarly, competition did not significantly modify the effect of global change drivers on diversity, nor did type of global change factor exert a significant effect (Figs. 2d and 3d, Table S6). Again, individual studies showed varying results. For example, in 2 cases, nutrient addition decreased diversity and competition mitigated this decrease (i.e., the negative effect of nutrients was smaller with more competition). In contrast, in 7 cases, nutrients decreased diversity and competition amplified this negative effect (i.e., the negative impact was greater with full competition; Fig. 2d; Table S2). In 2 cases, warming (1) and increased precipitation (1) increased richness and competition mitigated this increase. There were, again, several neutral cases (Table S2) and no studies manipulating CO_2_ and drought.

### (4) Pathogens

***Biomass*** Nutrient addition both alone (3 cases) and together with warming (1 case) increased community biomass and pathogens mitigated this increase (Fig. 4a, Table S3). There were also 2 neutral cases (1 nutrients, 1 temperature).

**Figure 4.**
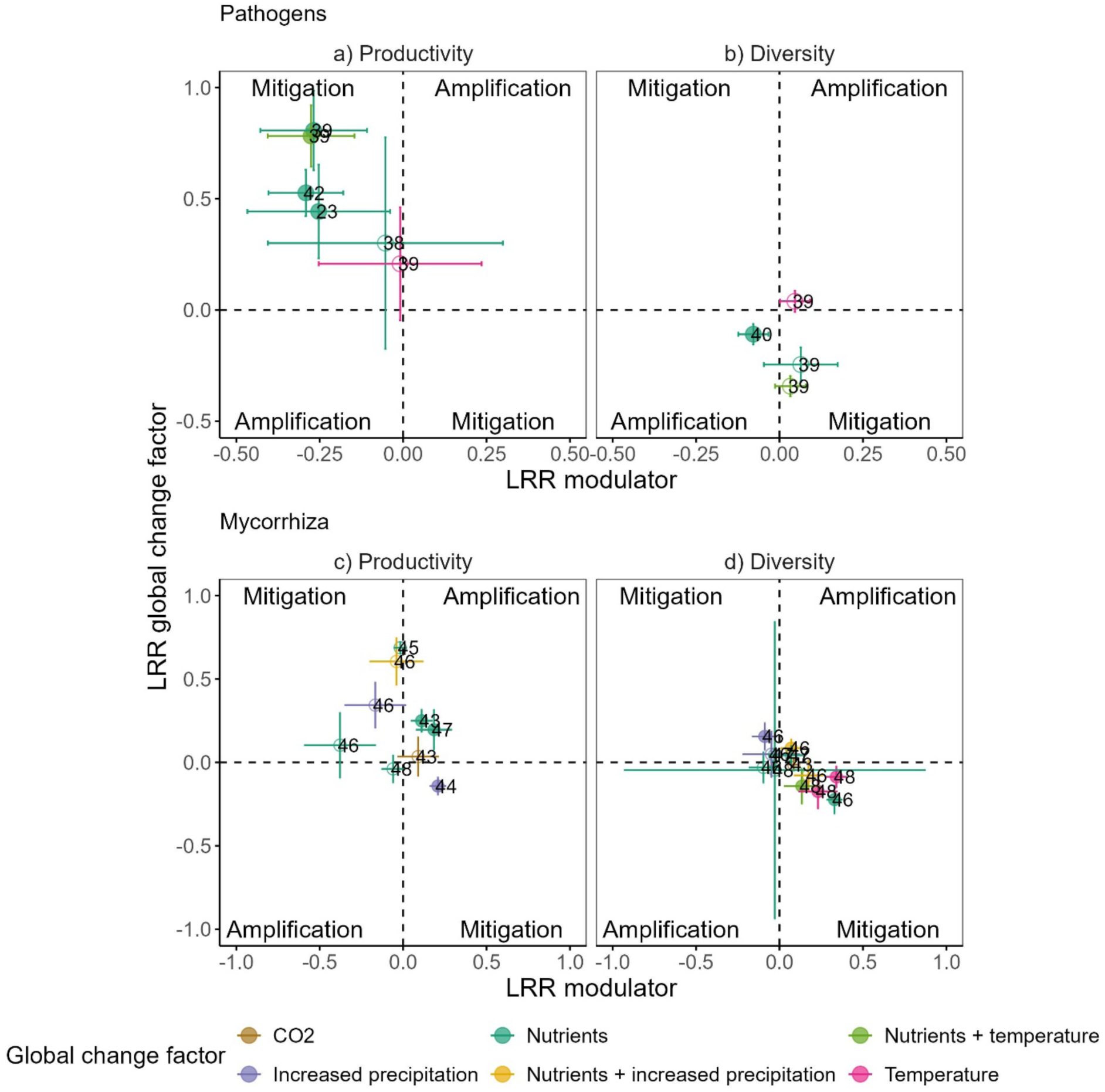
Dependence of the log response ratio (LRR) of diversity (a,c) and productivity (b,d) for the different global change drivers (increased precipitation, drought, nutrient increase, temperature, nutrients + temperature, nutrients + increased precipitation) and the LRR of the biotic modulators pathogens (a,b) and mycorrhiza (c,d). Positive values for LRR of the global change factor (y-axis) indicate a positive effect of the global change on diversity (a,c) or productivity (b,d) in the absence of the biotic modulators pathogens or mycorrhiza relative to the control in the absence of pathogens or mycorrhiza, whereas negative values indicate a negative effect of the global change on diversity (a,c) or productivity (b,d) in the absence of the biotic modulators pathogens or mycorrhiza relative to the control. As the LRR of the biotic modulator describes the impact of the global change factor in the absence relative to the presence of the modulators pathogens or mycorrhiza, mitigation (make the impact smaller) or amplification (make the impact greater) could be either indicated by positive or negative LRR values as it depends on the effect of the global change factor on diversity (a,c) and productivity (b,d). Different corners of the diagram indicate the direction of biotic modulation (mitigation, amplification). Filled circles indicate that the LRR and their SE do not overlap with zero, whereas open circles indicate an overlap with zero, i.e., neutral effect.

***Diversity*** Nutrient addition decreased diversity and pathogens amplified this decrease in 1 case. In 2 cases, nutrients alone or nutrients together with warming decreased plant diversity but pathogens had a neutral effect; however, there was tendency for mitigation in these cases (Fig. 4b, Table S3). There were no precipitation increase, drought or CO_2_ increase studies.

### (5) Mycorrhiza

***Biomass*** Nutrients increased biomass and mycorrhiza amplified this increase in 2 cases (Fig. 4c, Table S4). In 1 case, precipitation increase decreased biomass and mycorrhiza mitigated this decrease (Fig. 4c). Other cases were classified as neutral based on the overlap of SEs with zero in Fig. 4c; however, in two cases there was a trend of nutrients and precipitation increase enhancing biomass and mycorrhiza mitigating this increase (Fig. 4c).

***Diversity*** Global change factor (nutrients, warming, nutrients combined with warming) decreased plant diversity and mycorrhiza mitigated this decrease in 4 cases (Fig. 4d, Table S4). In 1 case, increased precipitation enhanced diversity and mycorrhiza mitigated this increment, while in 1 case, nutrients together with increased precipitation enhanced diversity and mycorrhiza amplified this increase. Other cases were classified as neutral (Fig. 4d, Table S4). There were no drought studies for mycorrhiza.

## IV. DISCUSSION

Our synthetic conceptual framework allowed us to integrate and compare how interspecific interactions, i.e., ‘biotic modulators’, alter global change impacts on plant community biomass and diversity. The strongest generalities emerging from synthesizing existing literature, testing our framework, were that herbivory significantly mitigated both enhanced plant biomass production and decline in plant diversity caused by different global change factors. The type of global change factor did not significantly affect our herbivory results, suggesting that our findings were uniform across global change factors. Our results were also concordant and strong across different terrestrial herbaceous ecosystems, ranging from low-productivity tundra grasslands to highly productive pampa meadows (Table S1). Further, the study systems were grazed by a variety of different wild and domestic herbivores, ranging from small mammals such as voles, lemmings and guinea pigs, to larger mammals such as reindeer, yak, and sheep (Table S1). Our findings extend and expand the prior individual studies and demonstrate that general patterns of herbivores mitigating global change effects do emerge across different global change drivers, herbivore types, and herbaceous systems.

### (1) Herbivores mitigate the impacts of global changes

Our finding that herbivory significantly mitigated biomass increment caused by global change factors, mainly by nutrient enrichment, is in line with earlier theoretical and empirical work showing that food-limited herbivores can consume a considerable proportion of aboveground productivity (Oksanen *et al*., 1981; Aunapuu *et al*., 2008; Jia *et al*., 2018; Borer *et al*., 2020; Trepel *et al*., 2024). Moreover, as a novel finding not shown in previous reviews or multisite global experiments (e.g., Borer *et al*., 2014), we show that herbivory across various herbivore guilds (including small and large vertebrates, and invertebrates) mitigated diversity decline caused by different global change factors, especially nutrient enrichment and climate warming. For example, in Borer et al. (2014), the authors found that both herbivores and nutrients independently influenced diversity through their effects on light limitation, but herbivory did not mitigate the negative effect of nutrient enrichment on diversity in this global grassland study. Most primary studies included in our meta-analysis did not measure both biomass and diversity; however, in those that did, mitigation of diversity decline by herbivores sometimes occurred even without any effect on biomass (Okach *et al*., 2019, Furey & Tilman, 2024), while in other cases both effects were found (Yang et al., 2015; Veen *et al*., 2024). Multiple mechanisms promoting plant coexistence can link herbivory to increased plant diversity along resource-gradients, some of which involve reduced biomass (e.g. alleviation of asymmetric light competition by consumption of tall and fast-growing species, growth-defense trade-off; Coley, Bryant & Chapin, 1985; Lind *et al*., 2013; Borer *et al*., 2014; Terborgh, 2015; Eskelinen *et al*., 2022), whereas others operate independently of biomass (e.g. reduction of litter, promotion of recruitment through creation of recruitment microsites and facilitating colonization from regional species pool (Olff & Ritchie, 1998; Eskelinen, Kaarlejärvi & Olofsson, 2017; Jessen *et al*., 2023). Our results give strong support for herbivores being important stabilizing agents that can help to combat global change effects on terrestrial plant communities in the Anthropocene.

Most primary studies manipulated nutrients, and we found only a few case studies addressing interactions among herbivores and climate. Among the few studies exploring interactions between warming and herbivory, although some reported relatively weak effects (Fig. 2; Moise & Henry, 2012; Wang *et al*., 2012; Kaarlejärvi, Eskelinen & Olofsson, 2013), some demonstrated that herbivory can mitigate climate warming effects on biomass and diversity (Post & Pedersen, 2008; Post *et al*., 2023) as well as the joint effects of warming and nutrient enrichment on diversity (Kaarlejärvi, Eskelinen & Olofsson, 2017). These experiments were mostly carried out in temperature-limited ecosystems where warming can both improve conditions for growth and also enhance soil nutrient availability through accelerating decomposition and nutrient cycling (Doetterl *et al*., 2022). Thus, warming can interact with herbivory similarly to how nutrient enrichment interacts with herbivory. Interestingly, while we were unable to test whether herbivory would differently modulate the effects of resource enhancing global change factors vs. resource-decreasing global change factors, one primary study that experimentally decreased water availability found that drought diminished biomass and herbivory amplified this negative effect (Okach *et al*., 2019). As demonstrated by this rare experimental study, we would expect grazing to exacerbate the effects of resource-diminishing global change factors.

### (2) Mixed effects of plant competition as a modulator

We found mixed effects of competition as a modulator (i.e., amplification, mitigation and neutral effects), and thus competition did not significantly alter global change effects on diversity and productivity overall in our cross-study meta-analysis. These heterogeneous responses may partially reflect varying methodology to manipulate competition (Table S2). However, in some primary studies competition mitigated the effects of nutrient enrichment on plant biomass (Hautier, Niklaus & Hector, 2009; Farrer & Suding, 2016), suggesting that competition can limit biomass production in nutrient-enriched conditions. This finding could reflect competition for light restraining the growth of species with the highest aboveground biomass production. Alternatively, it could indicate that when competition for light is strongest and communities are dominated by tall species superior in competition for light, the simultaneous loss of subordinate and small species significantly reduces total community biomass (Hautier, Niklaus & Hector, 2009), e.g., via reduced complementarity. Contrary to these findings, other primary studies reported that competition amplified biomass increment (Chaves & Smith, 2021), suggesting that the absence of dominant species can limit the response of total community biomass to global change factors. However, these responses may represent an artefact of the experimental design as removing a highly productive dominant species may inevitably diminish total community biomass.

We found several primary studies where nutrient enrichment decreased diversity and competition amplified this decline (Eskelinen 2010; Dickson & Foster, 2011; Dickson *et al*., 2014; Li *et al*., 2015; Doležal *et al*., 2019; Eskelinen *et al*., 2022). In some cases, although often not measured in the same study, the presence of highly competitive species amplified both biomass increase and diversity decline (Dickson *et al*., 2014), demonstrating how altered competitive interactions modify the effects of resource enhancing global change factors on diversity. It is noteworthy, however, that nutrient enrichment can also diminish diversity without changing biomass through reducing the dimensionality of belowground nutrient trade-offs (Harpole *et al*., 2016) or toxicity of nutrient effects (Britto & Kronzucker, 2002).

Very little is known about how competition interacts with global change factors that reduce resource availability (drought) or that affect resource availability indirectly (warming). For example, we found no experimental studies addressing how competition modifies drought effects on plant community diversity and productivity. This lack of studies likely stems from that water as a resource can be reduced through both abiotic (e.g., evaporation) and biotic (competition) means, which makes investigating its role challenging (Craine & Dybzinski, 2013). The only experimental study addressing interaction between warming and competition captured by our literature survey was conducted in tundra and found that warming increased seedling richness more in the absence of adult competitors than in their presence (Eskelinen, Kaarlejärvi & Olofsson, 2017). The authors demonstrated that competition can limit positive diversity response to climate warming, at least in early life stages. Taken together, while plant competition can interact with drought and warming to impact diversity and productivity, a paucity of studies poses a major gap in our understanding of these relationships.

### (3) Pathogens as modulators tend to behave similarly to herbivores

Although understanding the shifting role of pathogens in response to global change drivers is of critical importance (Paseka *et al*., 2020), data was extremely scarce. Some of the primary studies that we reviewed indicate that foliar pathogens can mitigate biomass enhancement caused by nutrient enrichment similarly than herbivores (Yan *et al*., 2023; Zaret *et al*., 2023; Zhang *et al*., 2024). These effects on biomass can arise from that nutrient enrichment can increase pathogens (Lekberg *et al*., 2021) through favoring fast-growing but less defended plants that are more susceptible to pathogens (i.e., growth-defense trade-off; Cappelli *et al*., 2020; He, Webster & He, 2022). In one primary study, foliar pathogens also tended to mitigate diversity loss caused by nutrient enrichment (although this was not significant based on our SE assessment; Yan *et al*., 2023). These findings are in line with earlier theoretical studies suggesting that pathogens can limit plant biomass and maintain plant diversity if their effects are density-dependent, and they attack and suppress the dominant plant species (Allan, Van Ruijven & Crawley, 2010; Bagchi *et al*., 2014). However, in another study, pathogens (both foliar fungal and root oomycetes) amplified the decline in diversity caused by nutrient enrichment (Zhao *et al*., 2023). Our findings suggest that pathogens may act relatively similarly to herbivores in terms of biomass, underscoring the broader role of consumers in stabilizing global change effects on plant communities.

### (4) Mycorrhiza can amplify or mitigate global changes effects

Similarly to pathogens, we found very few studies addressing the joint effects of mycorrhiza and global changes on plant community biomass and diversity. In studies captured by our literature survey, mycorrhiza tented to either amplify (Johnson, Wolf & Koch, 2003; Kang *et al*., 2020) or had a neutral impact (Van Der Heijden, Verkade & De Bruin, 2008) on biomass increment under nutrient enrichment, and in one case nutrient enrichment decreased biomass only in the presence of mycorrhiza (Xang *et al*., 2021). These findings cautiously suggest that mycorrhiza can exacerbate nutrient enrichment effects on community productivity. As soil fertility can dictate the cost and benefits from the symbiotic plant-mycorrhizal relationship (Johnson *et al*., 2015), mycorrhizal modification of plant community biomass can be complex; however, these responses may reflect, for example, enhanced nutrient interception and soil water retention, or prevention of leaching (Cavagnaro *et al*., 2015; Martínez-García *et al*., 2017) that could all amplify nutrient enrichment effects. However, Martinez-Gracia *et al*. (2017) found that community biomass slightly decreased due to increased precipitation and the presence of mycorrhiza mitigated this decrease; this finding demonstrates that mycorrhiza can also stabilize biomass production in response altered rainfall. For diversity we discovered that mycorrhiza mitigated the negative impacts of both nutrient enrichment (Yang *et al*., 2021a) and warming (Yang *et al*., 2021b) on diversity, further highlighting the stabilizing effects of mycorrhiza. These effects could reflect, for example, redistribution of nutrients and water through mycorrhizal networks between dominant and subdominant species (Van Der Heijden *et al*., 1998; Yang *et al*., 2018). Overall, these results suggest that mycorrhiza can alter the effects of different global change factors on plant community biomass and diversity.

### (5) Methodological challenges, knowledge gaps, and future directions

One of the most striking findings from our synthesis was the dearth of experimental studies testing the role of biotic modulators at the plant community level. For example, studies that addressed the role of plant competition, mycorrhiza, pathogens, and insect herbivores rarely reported treatment effects on whole community biomass and diversity but rather at the individual level, which cannot be used to infer responses of total community biomass and diversity. This likely reflects methodological challenges in conducting manipulations on community level (e.g., how to manipulate competition on community level) and in the field (e.g., how to manipulate mycorrhiza and insect herbivores in the field). For example, in competition studies, removing a highly productive dominant species may inevitably affect productivity, and removing a different species might cause distinct responses. In field studies, fungicides used to manipulate mycorrhiza in plant roots or soil pathogens can also alter other fungi living in the soil. Random biomass/species removals or manipulations that do not alter the physiological environment (such as light addition; Hautier, Niklaus & Hector, 2009; Eskelinen *et al*., 2022), would be better options. We advocate developing more unified methodology, reporting results on community level, and developing new technology, that directly addresses the role of biotic modulators, and would therefore considerably advance understanding the role of biotic modulators in mitigating/amplifying global change effects.

Very few studies manipulated both a biotic modulator and either drought, precipitation increase, or CO_2_ enrichment. For example, we know very little about herbivore impact on plant biomass and diversity in dryland ecosystems experiencing high warming rates and water shortages, although a negative impact of herbivory on plant biomass and diversity would be expected in these systems experiencing increasing pressure of livestock grazing (Maestre *et al*., 2022). Moreover, none of the warming studies were carried out in high-temperature systems, where warming is connected to increasing evapotranspiration and harsher, drying climate, and could interact with biotic modulators the opposite way than in temperature-limited, cold ecosystems (Harrison, 2020).

Finally, neither global changes nor biotic interactions happen in isolation from each other. For example, simultaneously acting global change drivers can alter plant communities in ways differing from those expected from single factor effects (Komatsu *et al*., 2019; Speißer *et al*., 2022). Similarly, although experiments addressing this issue are currently missing, multiple biotic factors (e.g., herbivory and pathogens) could synergize or antagonize to impact each other’s effects. It is also noteworthy that, theoretically, some (but not all) of the biotic modulation by herbivores and pathogens should operate through their effects on plant competition (Holt, Grover & Tilman, 1994; Huisman *et al*., 1999; Bradley, Gilbert & Martiny, 2008; Bever, Mangan & Alexander, 2015). Thus, one major avenue for future research would be to construct experiments that manipulate multiple factors, isolating changing conditions and multitrophic interactions, and direct and indirect pathways of their effects.

## V. CONCLUSIONS

1. Our conceptual framework of biotic modulators provides a comprehensive overview of how and by which mechanisms biotic interactions can modify global change effects on plant biodiversity and biomass (Fig. 1). This knowledge can be used for preditive hypothesis testing to uncover why local community responses to global changes are highly variable.
2. Our synthesis unravels remarkably few studies that can directly address causal biotic modulation at the plant community level. There is an urgent need for well-designed experiments addressing this research gap.
3. Several recent reviews highlight the role of herbivores, for example, in controlling alien plant proliferation (Mungi *et al*., 2023), modifying ecosystem properties and spatial heterogeneity (Trepel *et al*., 2024), and enhancing ecosystem carbon capture and storage (Schmitz *et al*., 2023). Here, our synthetic review expands these previous findings by providing quantitative evidence of herbivory as an effective nature-based control agent that can halt or decelerate plant biomass enhancement and biodiversity loss under changing environmental and climatic conditions. We allege that herbivory, in right intensity and frequency and in right places, can be used to prevent and restore global change effects on plant biodiversity and biomass. Moreover, although the data for pathogens was very scarce, our review implies that, in natural communities, pathogens may also act as nature’s own conservation agents that can counteract human-induced plant biomass enhancement, similarly to herbivores. Together, our results underscore the broader role of consumers in stabilizing plant community responses to global change.
4. Although we found mixed effects of plant competition as a modulator of global change effects, in several primary studies competition amplified nutrient enrichment caused plant biodiversity loss, highlighting plant competition as a mechanism leading to biodiversity loss. Our scarce data for mycorrhiza also shows that mycorrhiza can both exacerbate or stabilize global changes effects.
5. Overall, our synthesis reveals that biotic interactions, either within or between trophic levels, can critically alter global change effects on plant community biomass and diversity. These results will not only help us comprehending how, why and where plant communities change, but also aid us to unravel how these changes can be prevented and restored.

## VI. ACKNOWLEDGEMENTS

This work was supported by the Research Council of Finland (projects no 297191 and 351089) to A.E. and by Oulu University Biodiverse Anthropocenes Research Program (ANTS) to A.E. and L.K. We thank Gabriele Rada (iDiv, Media and Communications) for support with the graphics in Fig. 1.

## VII. AUTHOR CONTRIBUTIONS

A.E. had the original idea for the study. A.E. and L.K. conceived the study and generated more specific ideas. W.S.H. helped in conceptualizing the paper and all other authors contributed to generating ideas for the study. L.K. and A.E. planned the literature survey and L.K. planned the meta-analyses. A.E. and L.K. designed Figure 1 with help from W.S.H. and L.K. designed other figures. M.A. performed the review and meta-analyses with help from L.K., and drew Figures 2-4. A.E. wrote the manuscript with contributions from L.K. and all other authors commented the manuscript.

## VIII. DATA AVAILABILITY STATEMENT

Data and code will be made accessible upon acceptance in Dryad repository.

## X. SUPPORTING INFORMATION

Additional supporting information may be found online in the Supporting Information section at the end of the article.

**Search phrases for each biotic modulator.** Search phrases used in the literature search for herbivory, plant competition, pathogens and mycorrhiza.

**Table S1.** Detailed information about primary studies included in quantitative meta-analyses with herbivory as a biotic modulator.

**Table S2.** Detailed information about primary studies included in quantitative meta-analyses with competition as a biotic modulator.

**Table S3.** Detailed information about primary studies found in systematic literature review with pathogens as a biotic modulator.

**Table S4.** Detailed information about primary studies found in systematic literature review with mycorrhiza as a biotic modulator.

**Figure S1.** Funnel plots of different combinations of herbivory and competition (modulators) and diversity and competition (response variables).

**List of studies included in the two meta-analyses and two reviews.**

**Table S5.** Results of linear mixed effects meta-regression models for diversity and productivity as response variables and herbivory as the biotic modulator.

**Table S6.** Results of linear mixed effects meta-regression models for diversity and productivity as response variables and competition as the biotic modulator.

**Figure S2.** Modified PRISMA diagram for herbivory.

**Figure S3.** Modified PRISMA diagram for competition.

**Figure S4.** Modified PRISMA diagram for pathogens.

**Figure S5.** Modified PRISMA diagram for mycorrhiza.

