## supplements for "Biotic modulators of global change effects on plant communities"

**Supplementary Information**

**Search phrases for each biotic modulator.**

***Herbivory***: ((((TS=((herbivor*OR grazing OR grazer* OR “herbivore exclusion” OR “herbivore removal” OR “herbivory exclusion” OR “herbivory removal” OR  “consumer removal” OR “consumer exclusion” OR fencing OR “grazer exclusion” OR “grazer removal” OR “grazer manipulation” OR “grazing exclusion” OR “grazing removal”  OR “grazing removal “ OR “insect exclusion” OR “insect removal” OR “invertebrate exclusion” OR “invertebrate removal”))) AND TS=((“species richness” OR “seedling richness” OR “seedling abundance” OR “seedling number” OR evenness OR dominance OR diversity OR composition OR biomass OR productivity OR ANPP) )) AND TS=((plant community OR plant* OR vegetation))) AND TS=((experiment OR manipulation OR treatment OR “global change” OR “climate change”) )) AND TS=((water OR drought OR irrig* OR rainfall OR precipitation OR temperature OR warm* OR “nutrient enrichment” OR “nutrient addition” OR “nitrogen enrichment” OR “nitrogen addition” OR “N addition” OR “NPK addition” OR “NPK fertiliz*” OR fertiliz* OR “nitrogen deposition” OR “nutrient deposition” OR “CO2 enrich*” OR  CO2 manipul* OR “CO2 addition” OR CO2)). This search yielded 447 results (December 12, 2023). After screening manually through the titles and abstracts and in some cases the methods/results, 26 papers were found to be relevant (Table S1).

***Plant competition***: ((((TS=( (competition OR “competitor removal” OR “competition removal” OR “competitor reduction” OR “competition reduction” OR “neighb* removal” OR disturbance OR “dominant removal” OR “competitor manipulation” OR “competition manipulation” OR “neighb* manipulation” OR “biomass removal” OR facilitation))) AND TS=((“species richness” OR “seedling richness” OR “seedling abundance” OR “seedling number” OR evenness OR dominance OR diversity OR composition OR biomass OR productivity OR ANPP) )) AND TS=((plant community OR plant* OR vegetation))) AND TS=((experiment OR manipulation OR treatment OR “global change” OR “climate change”) )) AND TS=((water OR drought OR irrig* OR rainfall OR precipitation OR temperature OR warm* OR “nutrient enrichment” OR “nutrient addition” OR “nitrogen enrichment” OR “nitrogen addition” OR “N addition” OR “NPK addition” OR “NPK fertiliz*” OR fertiliz* OR “nitrogen deposition” OR “nutrient deposition” OR “CO2 enrich*” OR  CO2 manipul* OR “CO2 addition” OR CO2)). This search yielded 5763 results (December 12, 2023). After screening manually through the titles and abstracts and in some cases the methods/results, 14 papers were found to be relevant (Table S2).

***Pathogens***: ((((TS=((pathogen* OR “fungal pathogen” OR “fungal exclusion” OR “pathogen exclusion” OR “pathogen removal” OR fungicide))) AND TS=((“species richness” OR “seedling richness” OR “seedling abundance” OR “seedling number” OR evenness OR dominance OR diversity OR composition OR biomass OR productivity OR ANPP))) AND TS=((plant community OR plant* OR vegetation))) AND TS=((experiment OR manipulation OR treatment OR “global change” OR “climate change”))) AND TS=((water OR drought OR irrig* OR rainfall OR precipitation OR temperature OR warm* OR “nutrient enrichment” OR “nutrient addition” OR “nitrogen enrichment” OR “nitrogen addition” OR “N addition” OR “NPK addition” OR “NPK fertiliz*” OR fertiliz* OR “nitrogen deposition” OR “nutrient deposition” OR “CO2 enrich*” OR CO2 manipul* OR “CO2 addition” OR CO2)). This search yielded 1818 results (December 12, 2023). After screening manually through the titles and abstracts and in some cases the methods/results, 5 papers were found to be relevant (Table S3).

***Mycorrhiza:*** ((((TS=((“mycorrhiz* manipulation” OR mycorrhiz*OR AMF OR AM OR “mycorrhiz* treatment” OR “mycorrhiz* inoculation” OR “AMF inoculation” OR “AM inoculation”))) AND TS=((“species richness” OR “seedling richness” OR “seedling abundance” OR “seedling number” OR evenness OR dominance OR diversity OR composition OR biomass OR productivity OR ANPP))) AND TS=((plant community OR plant* OR vegetation))) AND TS=((experiment OR manipulation OR treatment OR “global change” OR “climate change”) )) AND TS=((water OR drought OR irrig* OR rainfall OR precipitation OR temperature OR warm* OR “nutrient enrichment” OR “nutrient addition” OR “nitrogen enrichment” OR “nitrogen addition” OR “N addition” OR “NPK addition” OR “NPK fertiliz*” OR fertiliz* OR “nitrogen deposition” OR “nutrient deposition” OR “CO2 enrich*” OR CO2 manipul* OR “CO2 addition” OR CO2)). This search yielded 761 results (December 12, 2023). After screening manually through the titles and abstracts and in some cases the methods/results, 6 papers were found to be relevant (Table S4).

**Table S1.** Primary studies included in quantitative meta-analyses with herbivory as a biotic modulator. Numbers in square brackets refer to study identities in Figs. 2a and c. System indicates the ecosystem/study system where the experiment was carried out. In some studies, multiple global change factors and/or several biotic modulators were manipulated. Global change outcome indicates the pure effect of global change factor in the absence of the modulator. In case of several global change factors, we separately report results for each global change factor and for their joint effect. Main finding refers to the direction of modulation; however, this direction does not indicate statistically significant differences. AGB, aboveground productivity; ANPP, aboveground net primary productivity; total biomass index, biomass estimate obtained using point intercept method; richness, species number.

| Study | System / Herbivores | Global change factors | Modulator treatment | Response variable | Main finding | Other modulators | Global change outcome |
|---|---|---|---|---|---|---|---|
| Okach et al. 2019 [1] | Humid savanna / Livestock | Increased precipitation | Vertebrate exclusion | Richness | mitigation | None | decrease |
| | | | | Shannon | mitigation | | decrease |
| | | | | AGB | amplification | | decrease |
| | | Drought | | Richness | mitigation | | decrease |
| | | | | Shannon | mitigation | | decrease |
| | | | | AGB | amplification | | decrease |
| Eskelinen et al. 2017 [2] | Tundra grassland / Reindeer, voles | Nutrients | Vertebrate exclusion | Richness | mitigation | Competition | decrease |
| | | Temperature | | Richness | amplification | | Increase |
| | | Nutrients + temperature | | Richness | mitigation | | decrease |
| Moise et al. 2012 [3] | Temperate old field grassland / Rodents and molluscs | Temperature | Insecticide | AGB | neutral | None | neutral |
| | | | Vertebrate exclusion | | neutral | | neutral |
| | | Nutrients | Insecticide | AGB | neutral | None | neutral |
| | | | Vertebrate exclusion | | mitigation | | increase |
| Lin et al. 2023 [4] | Temperate grassland / Livestock | Increased precipitation | Vertebrate exclusion | Richness | neutral | None | increase |
| | | | | Shannon | neutral | | decrease |
| | | | | AGB | mitigation | | increase |
| | | Drought | | Richness | mitigation | | decrease |
| | | | | Shannon | neutral | | decrease |
| | | | | AGB | neutral | | neutral |
| La Pierre et al. 2014 [5] | Prairie / Insects, mice and voles | Nutrients | Vertebrate exclusion | AGB | neutral | None | increase |
| | | | Insecticide | AGB | neutral | | increase |
| | | | Vertebrate exclusion | Richness | mitigation | | decrease |
| | | | Insecticide | Richness | neutral | | decrease |
| Eskelinen 2010 [6] | Non-acidic heath / Reindeer, voles | Nutrients | Vertebrate exclusion | Seedling richness | neutral | Competition | decrease |
| | Acidic heath / Reindeer, voles | | | Seedling richness | neutral | | neutral |
| Campana et al. 2021 [7] | Humid pampa meadow / Cattle | Nutrients | Vertebrate exclusion | AGB | neutral | None | neutral |
| Borer et al. 2020 [8] | Old field grasslands / Multiple herbivores | Nutrients | Vertebrate exclusion | AGB | neutral | None | increase |
| Yang et al. 2015 [9] | Alpine grassland / Sheep, yak | Nutrients | Vertebrate exclusion | Richness | mitigation | None | decrease |
| | | | | AGB | mitigation | | increase |
| Kaarlejärvi et al. 2013 [10] | Tundra grassland / Reindeer, voles | Temperature | Vertebrate exclusion | AGB | neutral | None | neutral |
| | | Nutrients | | AGB | mitigation | | increase |
| | | Nutrients + Temperature | | AGB | mitigation | | increase |
| Borgström et al. 2016 [11] | Mesocosm / Grasshoppers | Nutrients | Enclosures with and without insects | AGB | neutral | None | neutral |

| | | | | | | | |
|---|---|---|---|---|---|---|---|
| Eskelinen et al. 2012 [12] | Non-acidic heath / Reindeer, voles | Nutrients | Vertebrate exclusion | Total biomass index | neutral | None | increase |
| | | | | Richness | mitigation | | decrease |
| | Acidic heath / Reindeer, voles | | | Total biomass index | mitigation | | increase |
| | | | | Richness | neutral | | decrease |
| Post et al. 2008 [13] | Tundra grassland / Caribou, muskoxen | Temperature | Vertebrate exclusion | Total biomass index | mitigation | None | increase |
| Post et al. 2023 [14] | Tundra grassland / Caribou, muskoxen | Temperature | Vertebrate exclusion | Simpson | mitigation | None | decrease |
| Kaarlejärvi et al. 2017 [15] | Tundra grassland / Reindeer, voles | Temperature | Vertebrate exclusion | Richness | neutral | None | neutral |
| | | Nutrients | | Richness | mitigation | | decrease |
| | | Nutrients + temperature | | Richness | neutral | | decrease |
| Furey & Tilman 2023 [16] | Old field grassland / Deer | Nutrients | Vertebrate exclusion | AGB | amplification | None | increase |
| | | | | Richness | mitigation | | decrease |
| Wang et al. 2012 [17] | Alpine meadow / Sheep | Temperature | Vertebrate enclosure | Richness | neutral | None | decrease |
| Alberti et al. 2011 [18] | Salt marsh steppe / Guinea pigs, rodents | Nutrients | Vertebrate exclusion | AGB | mitigation | None | increase |
| | | | | Richness | neutral | | decrease |
| Gough et al. 2012 [19] | Dry heath tundra / Caribou, voles | Nutrients | Vertebrate exclusion | ANPP | neutral | None | increase |
| | Moist acidic tundra / Caribou, voles | | | ANPP | amplification | | increase |
| Gough & Johnson 2017 [20] | Dry heath tundra / Caribou, voles | Nutrients | Vertebrate exclusion | Richness | neutral | None | decrease |
| | Moist acidic tundra / Caribou, voles | | | Richness | mitigation | | decrease |
| Veen et al. 2024 [21] | Semi-natural grassland / Deer, wild boar | Nutrients | Vertebrate exclusion | Biomass | mitigation | None | increase |
| | | | | Shannon | mitigation | | decrease |
| Jessen et al. 2023 [22] | Central European grassland / Sheep | Nutrients | Vertebrate exclusion | Seedling richness | neutral | Competition | neutral |
| Zaret et al. 2023 [23] | Old field grassland / Insects, deer | Nutrients | Vertebrate exclusion | Biomass | neutral | None | increase |
| | | | Insecticide | Biomass | mitigation | | increase |
| Schädler et al. 2008 [24] | Mesocosm / Grasshoppers and snails | Nutrients | Invertebrate enclosures | Richness | neutral | None | decrease |
| | | | | Biomass | neutral | | increase |
| Denyer et al. 2007 [25] | Chalk grassland / Rabbits | Nutrients | Vertebrate exclusion | Biomass | mitigation | None | increase |
| Blue et al. 2011 [26] | Old field grassland / Insects | Nutrients | Insecticide | Biomass | neutral | None | increase |

**Table S2.** Primary studies included in quantitative meta-analyses with competition as a biotic modulator. Numbers in square brackets refer to study identities in Figs. 2c and d. System indicates the ecosystem/study system where the experiment was carried out. In some studies, multiple global change factors and/or several biotic modulators were manipulated. Global change outcome indicates the pure effect of global change factor in the absence of the modulator. In case of several global change factors, we separately report results for each global change factor and for their joint effect. Main finding refers to the direction of modulation; however, this direction does not indicate statistically significant differences. AGB, aboveground productivity; ANPP, aboveground net primary productivity; total biomass index, biomass estimate obtained using point intercept method; richness, species number.

| Study | System | Global change factors | Modulator treatment | Response variable | Main finding | Other modulators | Global change outcome |
|---|---|---|---|---|---|---|---|
| Li et al. 2015 [27] | Alpine meadow | Nutrients | Dominant species removal | Richness | neutral | None | decrease |
| | | | | Shannon | amplification | | decrease |
| | | | | AGB | mitigation | | decrease |
| Li et al. 2018 [28] | Alpine meadow | Nutrients | Functional group (grass) removal | AGB | neutral | None | neutral |
| | | | | Richness | mitigation | | decrease |
| | | | Functional group (forb) removal | AGB | neutral | | neutral |
| | | | | Richness | neutral | | neutral |
| Eskelinen 2010 [6] | Non-acidic heaths | Nutrients | Biomass removal | Seedling richness | amplification | Herbivory | decrease |
| | Acidic heaths | | | Seedling richness | neutral | | neutral |
| Chaves et al. 2021 [29] | Tall-grass prairie | Increased precipitation | Dominant species removal | AGB | amplification | None | increase |
| | | Nutrients | | AGB | neutral | | neutral |
| | | Nutrients + increased precipitation | | AGB | neutral | | neutral |
| | | Increased precipitation | | Richness | mitigation | | increase |
| | | Nutrients | | Richness | neutral | | increase |
| | | Nutrients + increased precipitation | | Richness | neutral | | increase |
| | | Increased precipitation | | Shannon | amplification | | increase |
| | | Nutrients | | Shannon | neutral | | neutral |
| | | Nutrients + increased precipitation | | Shannon | neutral | | increase |
| Bret-Harte et al. 2008 [30] | Moist acidic tundra | Nutrients | Dominant species removal | Biomass | neutral | None | neutral |
| Farrer et al. 2016 [31] | Tall grass prairie | Nutrients | Low planting density | Biomass | mitigation | None | increase |
| | Alpine tundra | | | Biomass | mitigation | | increase |
| | Desert grassland | | | Biomass | mitigation | | increase |
| | Tall grass prairie | | | Shannon | neutral | | neutral |
| | Alpine tundra | | | Shannon | mitigation | | decrease |
| | Desert grassland | | | Shannon | neutral | | neutral |
| Leps 1999 [32] | Wet meadow | Nutrients | Dominant species removal | Biomass | neutral | None | increase |
| | | | | Richness | neutral | | decrease |
| Dickson et al. 2011 [33] | Old field grassland | Nutrients | Tie back light manipulation | Richness | amplification | None | decrease |
| | | | Clipping light manipulation | Richness | amplification | | decrease |
| Dolezal et al. 2019 [34] | Meadow | Nutrients | Dominant species removal | Richness | amplification | None | decrease |
| | | | | Biomass | neutral | | neutral |
| Eskelinen et al. 2022 [35] | Central European grassland | Nutrients | Light addition | Richness | amplification | None | decrease |
| | | | | Shannon | neutral | | neutral |
| Eskelinen et al. 2016 [2] | Tundra grassland | Nutrients | Biomass removal | Richness | neutral | Herbivory | neutral |
| | | Temperature | | Richness | mitigation | | increase |
| | | Nutrients + temperature | | Richness | neutral | | decrease |

| | | | | | | | |
|---|---|---|---|---|---|---|---|
| Dickson et al. 2014 [36] | Low-productivity grassland | Nutrients | Clonal species removal | Richness | amplification | None | decrease |
| | | | | Biomass | neutral | | increase |
| Jessen et al. 2023 [22] | Central European grassland | Nutrients | Light addition | Seedling richness | neutral | Herbivory | neutral |
| Hautier et al. 2009 [37] | Mesocosm | Nutrients | Light addition | AGB | mitigation | None | increase |

**Table S3**. Primary studies found in systematic literature review with pathogens as a biotic modulator. Numbers in square brackets refer to study identities in Figs. 4a and b. System indicates the ecosystem/study system where the experiment was carried out. In some studies, multiple global change factors and/or several biotic modulators were manipulated. Global change outcome indicates the pure effect of global change factor in the absence of the modulator. In case of several global change factors, we separately report results for each global change factor and for their joint effect. Main finding refers to the direction of modulation; however, this direction does not indicate statistically significant differences. AGB, aboveground productivity; ANPP, aboveground net primary productivity; total biomass index, biomass estimate obtained using point intercept method; richness, species number.

| Study | System | Global change factors | Modulator treatment | Response variable | Main finding | Other modulators | Global change outcome |
|---|---|---|---|---|---|---|---|
| Haugwitz et al. 2011 [38] | High altitude heath | Nutrients | Fungicide | AGB | neutral | None | neutral |
| Yan et al. 2022 [39] | Alpine meadow | Nutrients | Fungicide | AGB | mitigation | None | increase |
| | | | | Richness | neutral | | decrease |
| | | Temperature | | AGB | neutral | | neutral |
| | | | | Richness | neutral | | neutral |
| | | Nutrients + temperature | | AGB | mitigation | | increase |
| | | | | Richness | neutral | | decrease |
| Zhao et al. 2023 [40] | Alpine meadow | Nutrients | Fungicide + oomyceticade | Richness | amplification | None | decrease |
| Zaret et al. 2023 [23] | Old field grassland | Nutrients | Fungicide | Biomass | mitigation | None | increase |
| Zhang et al. 2024 [42] | Alpine meadow | Nutrients | Fungicide | Biomass | mitigation | None | increase |

**Table S4.** Primary studies included in quantitative meta-analyses with mycorrhiza as a biotic modulator. Numbers in square brackets refer to study identities in Figs. 4c and d. System indicates the ecosystem/study system where the experiment was carried out. In some studies, multiple global change factors and/or several biotic modulators were manipulated. Global change outcome indicates the pure effect of global change factor in the absence of the modulator. In case of several global change factors, we separately report results for each global change factor and for their joint effect. Main finding refers to the direction of modulation; however, this direction does not indicate statistically significant differences. AGB, aboveground productivity; ANPP, aboveground net primary productivity; total biomass index, biomass estimate obtained using point intercept method; richness, species number.

| Study | System | Global change factors | Modulator treatment | Response variable | Main finding | Other modulators | Global change outcome |
|---|---|---|---|---|---|---|---|
| Johnson et al. 2003 [43] | Mesocosm | Nutrients | Mesocosms with and without AM | Dry shoot biomass | amplification | None | increase |
| | | $CO_2$ | | Dry shoot biomass | neutral | | neutral |
| | | | | Richness | neutral | | neutral |
| Martinez-Garcia et al. 2017 [44] | Mesocosm | Increased precipitation | Mesocosms with and without AM | AGB | mitigation | None | decrease |
| Van der Heijden et al. 2008 [45] | Mesocosm | Nutrients | Mesocosms with and without AM | AGB | neutral | None | increase |
| Yang et al. 2021a [46] | Alpine meadow | Nutrients | Benomyl fungicide to reduce AM | AGB | neutral | None | neutral |
| | | Increased precipitation | | AGB | neutral | | increase |
| | | Nutrients + increased precipitation | | AGB | neutral | | increase |
| | | Nutrients | | Shannon | neutral | | neutral |
| | | Increased precipitation | | Shannon | mitigation | | increase |
| | | Nutrients + increased precipitation | | Shannon | amplification | | increase |
| | | Nutrients | | Richness | mitigation | | decrease |
| | | Increased precipitation | | Richness | neutral | | neutral |
| | | Nutrients + increased precipitation | | Richness | neutral | | neutral |
| Kang et al. 2019 [47] | Meadow steppe | Nutrients | Benomyl fungicide to reduce AM | AGB | amplification | None | increase |
| | | | | Richness | neutral | | neutral |
| | | | | Shannon | neutral | | neutral |
| Yang et al. 2021b [48] | Desert steppe | Temperature | Benomyl fungicide to reduce AM | Richness | mitigation | None | decrease |
| | | Nutrients | | Community productivity | neutral | | neutral |
| | | Temperature | | Simpson | mitigation | | decrease |
| | | Nutrients | | Simpson | neutral | | neutral |
| | | Nutrients + Temperature | | Simpson | mitigation | | decrease |

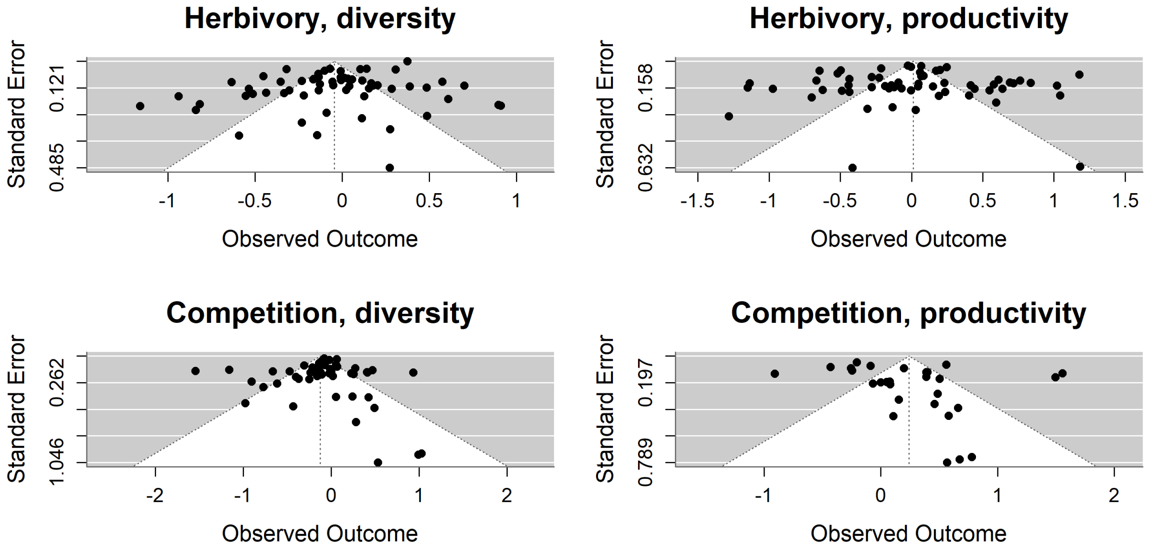


**Figure S1**. Funnel plots of different combinations of herbivory and competition (modulators) and diversity and competition (response variables). The effect size (observed outcome) of each study and response variable (dots) is plotted against their precision, measured as standard error. Dots outside the dashed lines can be an indication of publication bias.

**List of studies included in the two meta-analyses (herbivory, competition) and two reviews (pathogens, mycorrhiza).**


Alberti, J., Canepuccia, A., Pascual, J., Pérez, C. & Iribarne, O. (2011) Joint control by rodent herbivory and nutrient availability of plant diversity in a salt marsh-salty steppe transition zone: Rodent herbivory and nutrient availability controls on plant diversity. *Journal of Vegetation Science* **22**, 216–224.

Blue, J.D., Souza, L., Classen, A.T., Schweitzer, J.A. & Sanders, N.J. (2011) The variable effects of soil nitrogen availability and insect herbivory on aboveground and belowground plant biomass in an old-field ecosystem. *Oecologia* **167**, 771–780.

Borer, E.T., Harpole, W.S., Adler, P.B., Arnillas, C.A., Bugalho, M.N., Cadotte, M.W., Caldeira, M.C., Campana, S., Dickman, C.R., Dickson, T.L., Donohue, I., Eskelinen, A., Firn, J.L., Graff, P., Gruner, D.S., et al. (2020) Nutrients cause grassland biomass to outpace herbivory. *Nature Communications* **11**, 6036.

Borgström, P., Strengbom, J., Marini, L., Viketoft, M. & Bommarco, R. (2017) Above- and belowground insect herbivory modifies the response of a grassland plant community to nitrogen eutrophication. *Ecology* **98**, 545–554.

Bret-Harte, M.S., Mack, M.C., Goldsmith, G.R., Sloan, D.B., DeMarco, J., Shaver, G.R., Ray, P.M., Biesinger, Z. & Chapin, F.S. (2008) Plant functional types do not predict biomass responses to removal and fertilization in Alaskan tussock tundra. *Journal of Ecology* **96**, 713–726.

Campana, S. & Yahdjian, L. (2021) Plant quality and primary productivity modulate plant biomass responses to the joint effects of grazing and fertilization in a mesic grassland. *Applied Vegetation Science* **24**, e12588.

Chaves, F.A. & Smith, M.D. (2021) Resources do not limit compensatory response of a tallgrass prairie plant community to the loss of a dominant species. *Journal of Ecology* **109**, 3617–3633.

Denyer, J.L., Hartley, S.E. & John, E.A. (2007) Small mammalian herbivore determines vegetation response to patchy nutrient inputs. *Oikos* **116**, 1186–1192.

Dickson, T.L. & Foster, B.L. (2011) Fertilization decreases plant biodiversity even when light is not limiting: Fertilization, light and plant biodiversity. *Ecology Letters* **14**, 380–388.

Dickson, T.L., Mittelbach, G.G., Reynolds, H.L. & Gross, K.L. (2014) Height and clonality traits determine plant community responses to fertilization. *Ecology* **95**, 2443–2452.

Doležal, J., Lanta, V., Mudrák, O. & Lepš, J. (2019) Seasonality promotes grassland diversity: Interactions with mowing, fertilization and removal of dominant species. *Journal of Ecology* **107**, 203–215.

Eskelinen, A. (2010) Resident functional composition mediates the impacts of nutrient enrichment and neighbour removal on plant immigration rates. *Journal of Ecology*

**98**, 540–550.

Eskelinen, A., Harpole, W.S., Jessen, M.-T., Virtanen, R. & Hautier, Y. (2022) Light competition drives herbivore and nutrient effects on plant diversity. *Nature* **611**, 301–305.

Eskelinen, A., Harrison, S. & Tuomi, M. (2012) Plant traits mediate consumer and nutrient control on plant community productivity and diversity. *Ecology* **93**, 2705–2718.

Eskelinen, A., Kaarlejärvi, E. & Olofsson, J. (2017) Herbivory and nutrient limitation protect warming tundra from lowland species' invasion and diversity loss. *Global Change Biology* **23**, 245–255.

Farrer, E.C. & Suding, K.N. (2016) Teasing apart plant community responses to N enrichment: the roles of resource limitation, competition and soil microbes. *Ecology Letters* **19**, 1287–1296.

Furey, G.N. & Tilman, D. (2024) Trade-offs between deer herbivory and nitrogen competition alter grassland forb composition. *Oecologia* **204**, 47–58.

Gough, L. & Johnson, D.R. (2018) Mammalian herbivory exacerbates plant community responses to long-term increased soil nutrients in two Alaskan tundra plant communities. *Arctic Science* **4**, 153–166.

Gough, L., Moore, J.C., Shaver, G.R., Simpson, R.T. & Johnson, D.R. (2012) Above- and belowground responses of arctic tundra ecosystems to altered soil nutrients and mammalian herbivory. *Ecology* **93**, 1683–1694.

Haugwitz, M.S., Michelsen, A. & Schmidt, I.K. (2011) Long-term microbial control of nutrient availability and plant biomass in a subarctic-alpine heath after addition of carbon, fertilizer and fungicide. *Soil Biology and Biochemistry* **43**, 179–187.

Hautier, Y., Niklaus, P.A. & Hector, A. (2009) Competition for Light Causes Plant Biodiversity Loss After Eutrophication. *Science* **324**, 636–638.

Jessen, M., Auge, H., Harpole, W.S. & Eskelinen, A. (2023) Litter accumulation, not light limitation, drives early plant recruitment. *Journal of Ecology* **111**, 1174–1187.

Johnson, N.C., Wolf, J. & Koch, G.W. (2003) Interactions among mycorrhizae, atmospheric $CO_2$ and soil N impact plant community composition. *Ecology Letters* **6**, 532–540.

Kaarlejärvi, E., Eskelinen, A. & Olofsson, J. (2013) Herbivory prevents positive responses of lowland plants to warmer and more fertile conditions at high altitudes. *Functional Ecology* **27**, 1244–1253.

Kaarlejärvi, E., Eskelinen, A. & Olofsson, J. (2017) Herbivores rescue diversity in warming tundra by modulating trait-dependent species losses and gains. *Nature Communications* **8**, 419.

Kang, F., Yang, B., Wujisiguleng, Yang, X., Wang, L., Guo, J., Sun, W., Zhang, Q. & Zhang, T. (2020) Arbuscular mycorrhizal fungi alleviate the negative effect of nitrogen deposition on ecosystem functions in meadow grassland. *Land Degradation & Development* **31**, 748–759.

La Pierre, K.J., Joern, A. & Smith, M.D. (2015) Invertebrate, not small vertebrate, herbivory interacts with nutrient availability to impact tallgrass prairie community composition and forb biomass. *Oikos* **124**, 842–850.

Lepš, J. (1999) Nutrient status, disturbance and competition: an experimental test of relationships in a wet meadow. *Journal of Vegetation Science* **10**, 219–230.

Li, W., Cheng, J., Yu, K., Epstein, H.E. & Du, G. (2015) Short-term responses of an alpine meadow community to removal of a dominant species along a fertilization gradient. *Journal of Plant Ecology* **8**, 513–522.

Li, W., Zhang, R., Liu, S., Li, W., Li, J., Zhou, H. & Knops, J.M.H. (2018) Effect of loss of plant functional group and simulated nitrogen deposition on subalpine ecosystem properties on the Tibetan Plateau. *Science of The Total Environment* **631–632**, 289–297.

Lin, X., Zhao, H., Zhang, S., Li, R., Li, X. & Wang, S. (2024) Impact of precipitation and grazing on recovery of plant vegetation in temperate grasslands. *Land Degradation & Development* **35**, 1178–1191.

Martínez-García, L.B., De Deyn, G.B., Pugnaire, F.I., Kothamasi, D. & Van Der Heijden, M.G.A. (2017) Symbiotic soil fungi enhance ecosystem resilience to climate change. *Global Change Biology* **23**, 5228–5236.

Moise, E.R.D. & Henry, H.A.L. (2012) Interactions of herbivore exclusion with warming and N addition in a grass-dominated temperate old field. *Oecologia* **169**, 1127–1136.

Okach, D.O., Ondier, J.O., Rambold, G., Tenhunen, J., Huwe, B., Jung, E.Y. & Otieno, D.O. (2019) Interaction of livestock grazing and rainfall manipulation enhances herbaceous species diversity and aboveground biomass in a humid savanna. *Journal of Plant Research* **132**, 345–358.

Post, E., Kaarlejärvi, E., Macias-Fauria, M., Watts, D.A., Bøving, P.S., Cahoon, S.M.P., Higgins, R.C., John, C., Kerby, J.T., Pedersen, C., Post, M. & Sullivan, P.F. (2023) Large herbivore diversity slows sea ice–associated decline in arctic tundra diversity. *Science* **380**, 1282–1287.

Post, E. & Pedersen, C. (2008) Opposing plant community responses to warming with and without herbivores. *Proceedings of the National Academy of Sciences* **105**, 12353–12358.

Schädler, M., Rottstock, T. & Brandl, R. (2008) Do nutrients and invertebrate herbivory interact in an artificial plant community? *Basic and Applied Ecology* **9**, 550–559.

Van Der Heijden, M.G.A., Verkade, S. & De Bruin, S.J. (2008) Mycorrhizal fungi reduce the negative effects of nitrogen enrichment on plant community structure in dune grassland. *Global Change Biology* **14**, 2626–2635.

Veen, G.F. (Ciska), Vermaat, A.T., Sitters, J. & Bakker, E.S. (2024) Vertebrate grazing can mitigate impacts of nutrient addition on plant diversity and insect abundance in a semi-natural grassland. *Oikos* **2024**, e10422.

Wang, S., Duan, J., Xu, G., Wang, Y., Zhang, Z., Rui, Y., Luo, C., Xu, B., Zhu, X., Chang, X., Cui, X., Niu, H., Zhao, X. & Wang, W. (2012) Effects of warming and grazing on soil N availability, species composition, and ANPP in an alpine meadow. *Ecology* **93**, 2365–2376.

Yan, X., Kohli, M., Wen, Y., Wang, X., Zhang, Y., Yang, F., Zhou, X., Du, G., Hu, S. & Guo, H. (2023) Nitrogen addition and warming modulate the pathogen impact on plant biomass by shifting intraspecific functional traits and reducing species richness. *Journal of Ecology* **111**, 509–524.

Yang, X., Mariotte, P., Guo, J., Hautier, Y. & Zhang, T. (2021a) Suppression of arbuscular mycorrhizal fungi decreases the temporal stability of community productivity under elevated temperature and nitrogen addition in a temperate meadow. *Science of The Total Environment* **762**, 143137.

Yang, X., Tian, H., Feng, J., Zang, J., Ji, B., Wang, Z. & Shen, Y. (2021b) Inter-annual precipitation variability alters the effects of soil resource enrichment and mycorrhizal suppression on plant communities in desert steppe. *Journal of Vegetation Science* **32**, e13082.

Yang, Z., Hautier, Y., Borer, E.T., Zhang, C. & Du, G. (2015) Abundance- and functional-based mechanisms of plant diversity loss with fertilization in the presence and absence of herbivores. *Oecologia* **179**, 261–270.

Zaret, M., Kinkel, L., Borer, E.T. & Seabloom, E.W. (2023) Soil nutrients cause threefold increase in pathogen and herbivore impacts on grassland plant biomass. *Journal of Ecology* **111**, 1629–1640.

Zhang, P., Huang, M., Chen, C., Hu, K., Ke, J., Liu, M., Xiao, Y. & Liu, X. (2024) Contrasting roles of fungal and oomycete pathogens in mediating nitrogen addition and winter grazing effects on biomass. *Ecology* **105**, e4254.

Zhao, Y., Liu, X., Wang, J., Nie, Y., Huang, M., Zhang, L., Xiao, Y., Zhang, Z. & Zhou, S. (2023) Fungal pathogens increase community temporal stability through species asynchrony regardless of nutrient fertilization. *Ecology* **104**, e4166.

**Table S5.** Results of linear mixed effects meta-regression models for diversity and productivity as response variables and herbivory as the biotic modulator. In both models, fixed predictor variables were the type of manipulation (sole effect of global change factor or the impact of modulator) and the type of global change factor (nutrient enrichment, climate warming, increased precipitation, drought, and CO2 increase), and study was included study as a random factor. The models were weighted by the corresponding sample variance of LRR to account for the variability in the results of each primary study. Significant P-values (≤ 0.05) are indicated in bold.

| | Diversity | | | | | | Productivity | | | | | |
|---|---|---|---|---|---|---|---|---|---|---|---|---|
| | Estimate | se | zval | pval | ci.lb | ci.ub | Estimate | se | zval | pval | ci.lb | ci.ub |
| Intercept | -0.28 | 0.10 | -2.67 | <0.01 | -0.48 | -0.07 | -0.04 | 0.17 | -0.24 | 0.81 | -0.38 | 0.30 |
| Increased precipitation | -0.02 | 0.13 | -0.18 | 0.86 | -0.29 | 0.24 | 0.19 | 0.24 | 0.80 | 0.43 | -0.28 | 0.66 |
| Nutrients | -0.09 | 0.11 | -0.78 | 0.44 | -0.30 | 0.13 | 0.54 | 0.18 | 3.02 | **<0.01** | 0.19 | 0.89 |
| Nutrients and Temperature | -0.05 | 0.17 | -0.32 | 0.75 | -0.39 | 0.28 | 0.62 | 0.29 | 2.12 | **<0.05** | 0.05 | 1.19 |
| Temperature | 0.05 | 0.13 | 0.39 | 0.69 | -0.21 | 0.32 | 0.19 | 0.21 | 0.90 | 0.37 | -0.22 | 0.60 |
| Modulator | 0.55 | 0.07 | 7.47 | **<.0001** | 0.40 | 0.69 | -0.75 | 0.09 | -8.23 | **<.0001** | -0.93 | -0.57 |

**Table S6.** Results of linear mixed effects meta-regression models for diversity and productivity as response variables and competition as the biotic modulator. In both models, fixed predictor variables were the type of manipulation (sole effect of global change factor or the impact of modulator) and the type of global change factor (nutrient enrichment, climate warming, increased precipitation, drought, and CO2 increase), and study was included study as a random factor. The models were weighted by the corresponding sample variance of LRR to account for the variability in the results of each primary study. Significant P-values (≤ 0.05) are indicated in bold.

| | Diversity | | | | | | Productivity | | | | | |
|---|---|---|---|---|---|---|---|---|---|---|---|---|
| | Estimate | se | zval | pval | ci.lb | ci.ub | Estimate | se | zval | pval | ci.lb | ci.ub |
| Intercept | 0.19 | 0.26 | 0.73 | 0.47 | -0.32 | 0.71 | 0.55 | 0.55 | 1.00 | 0.32 | -0.53 | 1.63 |
| Nutrients | -0.40 | 0.26 | -1.53 | 0.13 | -0.92 | 0.11 | -0.34 | 0.54 | -0.64 | 0.52 | -1.41 | 0.71 |
| Temperature | -0.14 | 0.38 | -0.37 | 0.71 | -0.89 | 0.61 | NA | NA | NA | NA | NA | NA |
| Nutrients and Temperature | -0.27 | 0.38 | -0.69 | 0.49 | -1.02 | 0.49 | NA | NA | NA | NA | NA | NA |
| Nutrients and increased precipitation | 0.04 | 0.36 | 0.11 | 0.91 | -0.66 | 0.74 | 0.01 | 0.77 | 0.02 | 0.99 | -1.50 | 1.53 |
| Modulator | 0.04 | 0.12 | 0.32 | 0.75 | -0.20 | 0.28 | 0.02 | 0.22 | 0.10 | 0.92 | -0.42 | 0.46 |

Herbivory

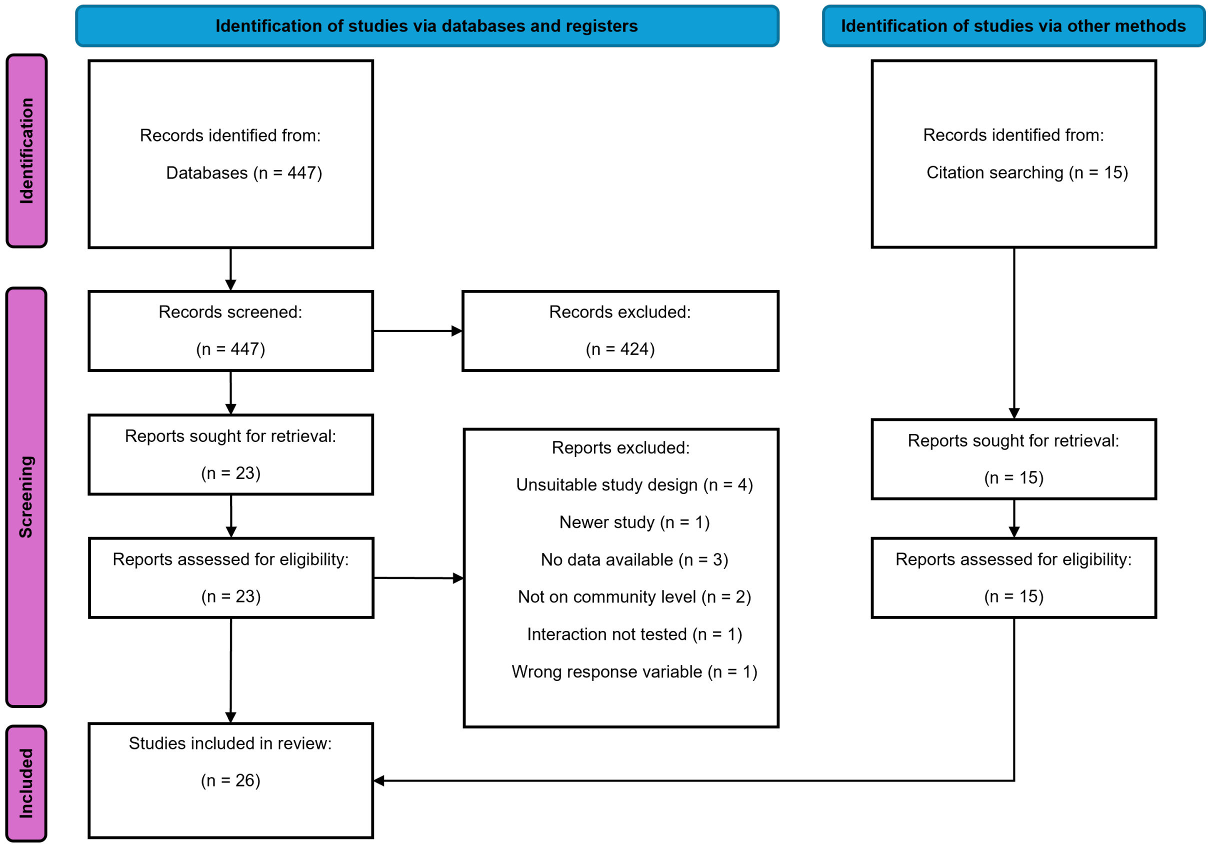


From: Page MJ, McKenzie JE, Bossuyt PM, Boutron I, Hoffmann TC, Mulrow CD, et al. The PRISMA2020 statement: an updated guideline for reporting systematic reviews. BMJ 2021;372:n71. doi: 10.1136/bmj.n71

**Figure S2**. Modified PRISMA diagram. Sequence of information in the different phases of the systematic review for herbivory as a modulator of global change effects on plant biomass and diversity following the PRISMA protocol and guidelines.

Competition

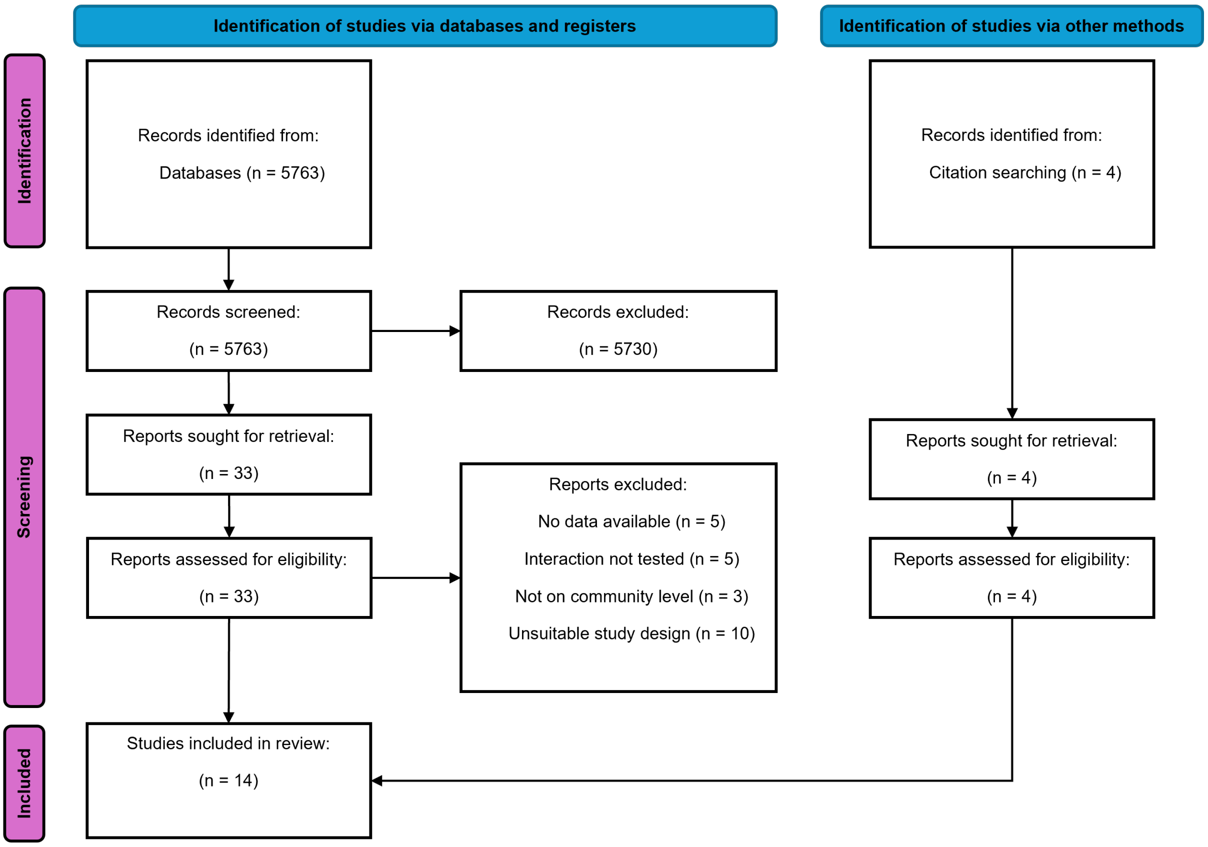


From: Page MJ, McKenzie JE, Bossuyt PM, Boutron I, Hoffmann TC, Mulrow CD, et al. The PRISMA2020 statement: an updated guideline for reporting systematic reviews. BMJ 2021;372:n71. doi: 10.1136/bmj.n71

**Figure S3**. Modified PRISMA diagram. Sequence of information in the different phases of the systematic review for competition as a modulator of global change effects on plant biomass and diversity following the PRISMA protocol and guidelines.

Pathogen

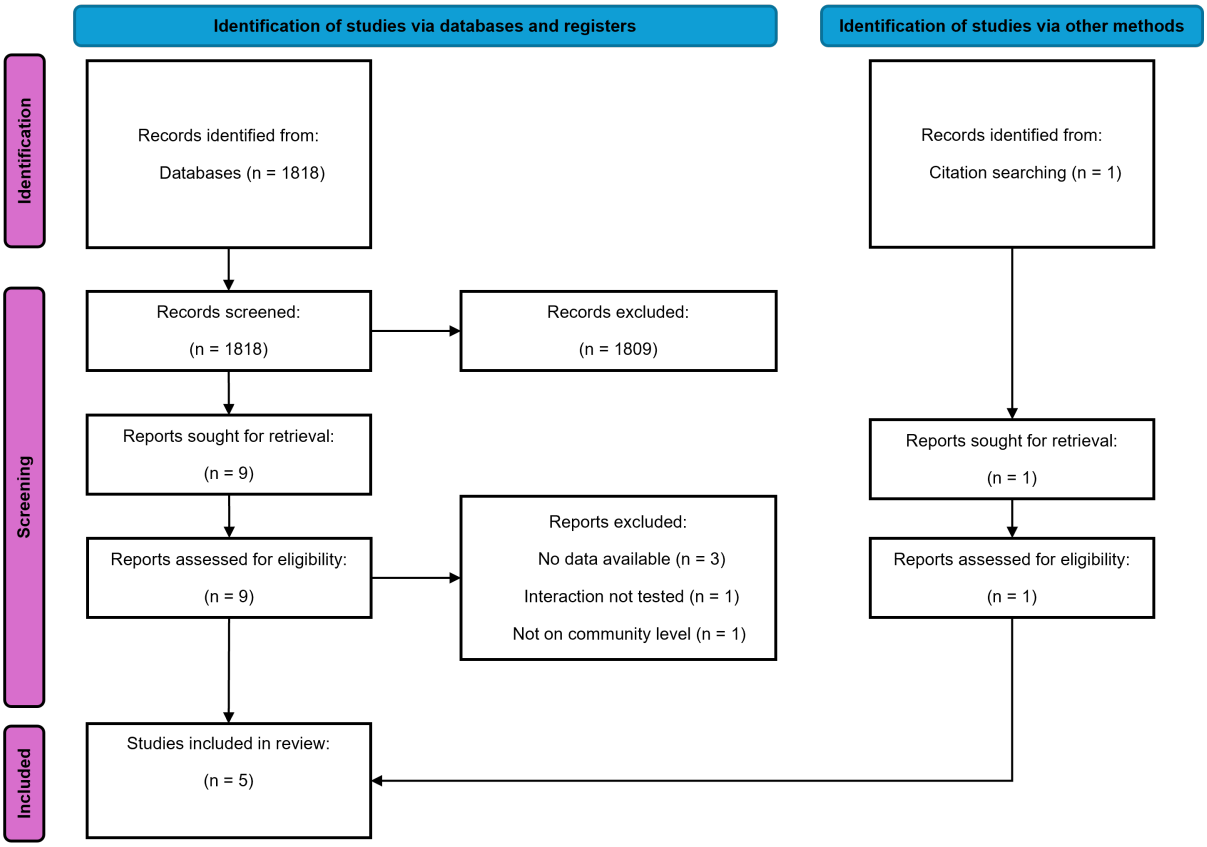


From: Page MJ, McKenzie JE, Bossuyt PM, Boutron I, Hoffmann TC, Mulrow CD, et al. The PRISMA2020 statement: an updated guideline for reporting systematic reviews. BMJ 2021;372:n71. doi: 10.1136/bmj.n71

**Figure S4**. Modified PRISMA diagram. Sequence of information in the different phases of the systematic review for pathogens as a modulator of global change effects on plant biomass and diversity following the PRISMA protocol and guidelines.

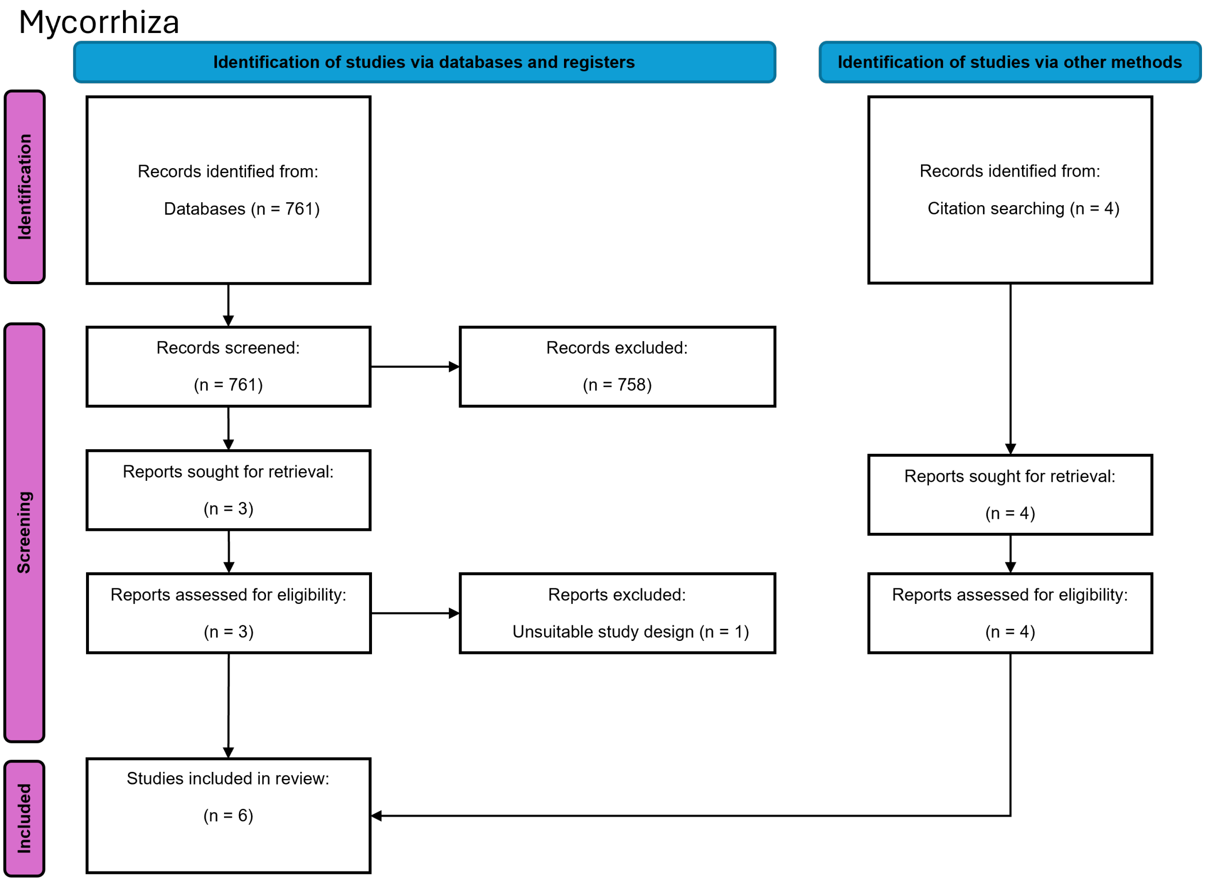


From: Page MJ, McKenzie JE, Bossuyt PM, Boutron I, Hoffmann TC, Mulrow CD, et al. The PRISMA2020 statement: an updated guideline for reporting systematic reviews. BMJ 2021;372:n71. doi: 10.1136/bmj.n71

**Figure S5**. Modified PRISMA diagram. Sequence of information in the different phases of the systematic review for mycorrhiza as a modulator of global change effects on plant biomass and diversity following the PRISMA protocol and guidelines.
